# Characterization of an Eye Field-like State during Optic Vesicle Organoid Development

**DOI:** 10.1101/2022.08.16.504119

**Authors:** Liusaidh J. Owen, Jacqueline Rainger, Hemant Bengani, Fiona Kilanowski, David R. FitzPatrick, Andrew S. Papanastasiou

## Abstract

Specification of the eye field (EF) within the neural plate marks the earliest detectable stage of eye development. Experimental evidence, primarily from non-mammalian model systems, indicates that the stable formation of this group of cells requires the activation of a set of key transcription factors (TFs). This critical event is challenging to probe in mammals and, quantitatively, little is known regarding the regulation of the transition of cells to this ocular fate. Using optic vesicle organoids to model the onset of the EF, we generate timecourse transcriptomic data allowing us to identify dynamic gene-expression programs that characterise this cellular-state transition. Integrating this with chromatin accessibility data suggests a direct role of canonical EFTFs in regulating these gene-expression changes, and highlights candidate *cis*-regulatory elements through which these TFs act. Finally, we begin to test a subset of these candidate enhancer elements, within the organoid system, by perturbing the underlying DNA sequence and measuring transcriptomic changes during EF activation.

## INTRODUCTION

Vertebrate eye development begins with the formation of the eye field – a patch of morphologically indistinct neuroectodermal cells within the anterior neural plate defined by co-expression of a set of transcription factors (TFs). The eye field was first identified in *Xenopus laevis* embryos (Zuber et al., 2003;Zuber, 2010) and expresses a specific set of TFs, known as the eye field transcription factors (EFTFs), with those being: *PAX6, RAX, SIX3, SIX6, LHX2, OTX2, NR2E1* and *TBX3* (using gene symbols of human orthologs). Mutations in at least three of the genes encoding orthologs of the *Xenopus* EFTF cause severe bilateral eye malformations in humans (*OTX2*, *PAX6* and *RAX*) (Fitzpatrick and van Heyningen, 2005). It seems likely that these genes are involved in eye field specification in humans. A better understanding of the genetic networks operating to form and maintain the eye field is important for both clinical genetics research and developmental biology.

A given cellular state can be thought of as the expression of a distinct combinatorial code of transcription factors, and differentiation – the transition from one cellular state to another – can be documented through the conceptual framework of Gene Regulatory Networks (GRNs) (Peter and Davison, 2011). A core concept in GRNs are *cis*-regulatory modules, sets of *cis*-regulatory elements (CREs – enhancers and repressors) that maintain the expression of the TF code for an individual cellular state as long as it is required. GRNs are established and maintained via cross- and auto-regulatory feedback loops with repression of TFs that are disruptive to the network. The EFTFs are thought to form part of a GRN essential for initiating eye development and the goal of this work is to characterise this GRN in mammals and gain a deeper understanding on how it is initiated and stably maintained.

The eye field has been difficult to study in mammals as it forms in early post-implantation embryos (~ 18 post ovulatory days in humans, ~ E7.5 in mouse). The Sasai lab developed a simple defined organoid culture system (Eiraku et al., 2011; Eiraku and Sasai, 2012; Nakano et al., 2012) that supports efficient differentiation of human and mouse embryonic stem cells (ESCs) into early eye structures; optic vesicles (OV) evaginate from the main body of the organoid and then develop into optic cup structures with partially differentiated neural retina. This transformative technology establishes that early eye development is a self-organising system and provides a window onto the hitherto cryptic processes driving the commitment to eye fate in mammals.

We have performed a functional genomic analysis of the eye field like cells that appear in the very early stages of optic vesicle organoid differentiation. By performing computational analyses of the transcriptomic changes during OV differentiation we find that the establishment of the eye field in mouse ESC-derived OVs, involves the strong up-regulation of a set of genes that includes the same key transcription factors identified in *Xenopus laevis*. Furthermore, by integrating transcriptomic and chromatin-accessibility data, we show that these TFs likely regulate the stable transition of cells to an ocular fate by binding distal enhancers in genomic domains local to EF genes. By examining how chromatin-accessibility, associated with key TFs, changes at individual genomic loci we generate hypotheses of plausible *cis*-regulatory elements required for the differentiation from pluripotency to eye field cell states. Finally, we test a subset of these hypotheses by perturbing the underlying DNA sequences. In all, by exploiting the robust and reproducible model system offered by the mESC-derived organoids, our study provides a first quantitative understanding of the regulatory mechanisms underlying mammalian eye field specification.

## RESULTS

### Mammalian EF genes overlap significantly with *Xenopus* EFTFs

Mammalian eye field specification – the crucial first stage of ocular development – is an event that occurs very early in development (during gastrulation), and as such is extremely challenging to study *in vivo*. We have exploited a reproducible, *in vitro* organoid model system, that closely mimics the *in vivo* regulatory dynamics, allowing us to generate data from this cell-state transition and through computational analysis gain a quantitative understanding of the underlying mechanisms. We performed OV organoid culture using mouse embryonic stem cells (mESCs) that have a GFP fluorescent reporter cassette knocked into the *Rax* locus (Eiraku and Sasai,2012) which expresses only in cells committed to an ocular fate. Rax-positive cells become detectable between day 3 and day 4 in culture (Fig.1A), and at days 5 and 6 ≳ 70%of live cells are Rax-positive (Table.S1). We performed bulk RNA-seq analysis using biological triplicates of OV organoid cultures at days 0, 1, 2, 3, 4 and 5. To enrich for cells in an eye field state, at day 4 and day 5 the cells were sorted using FACS into GFP positive and GFP negative populations (Fig.1B).

**Fig. 1.**
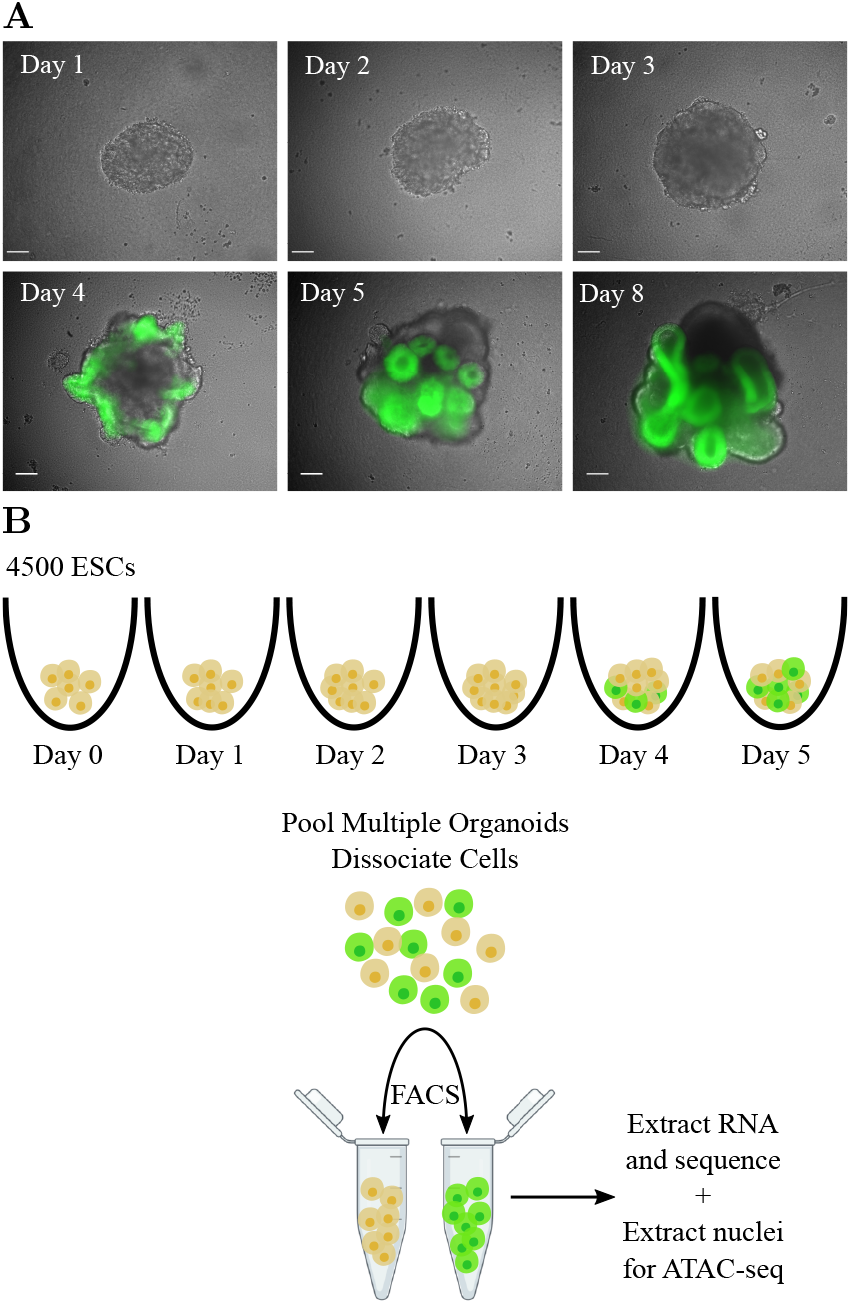
Exploiting optic vesicle organoids to probe the molecular origins of eye field specification. **A.** Representative organoids from timecourse days 1–5, and day 8. Scale bars: 100 *μ*m. **B.** Schematic of the experimental setup for timecourse RNA-seq and ATAC-seq data generation.

To identify genes characterizing the transition to the eye field state, we looked for differentially expressed genes between day 3 and the GFP-positive samples on days 4 and 5 (using `DEseq` (Love et al., 2014), FDR < 0.001, absolute log2 fold-change > 1.5). This analysis revealed a small set of “EF-up” genes (37) that are significantly up-regulated between these time points, of which 11 correspond to known mouse TFs (Fig.2A). Importantly, these include the canonical eye field genes Rax, *Pax6, Lhx2, Six3* and *Six6*, indicating that drivers of the optic-fate in mammals include most of the key TFs also discovered in *Xenopus*. Whist *Rax*, *Pax6*, *Lhx2* and *Six3* appear to be turned on between days 3 and 4, *Six6* appears to be expressed slightly later, consistent with evidence that it is a downstream target of *Lhx2* and *Pax6*(Tétreault et al., 2009). While not sufficiently up-regulated to pass our differential-expression threshold, *Nr2e1* displays an expression pattern consistent with that of the other canonical EFTFs. The only canonical EFTF with a somewhat unexpected expressionpattern – one that is anti-correlated with the onset of the eye-field – is *Tbx3*. This could be because *Tbx3* plays a different role in mouse than in *Xenopus* (*Tbx3* is a known component of the pluripotency network in mESCs (Russell, 2015)) or because the *Xenopus Tbx3* ortholog in mouse may not actually be *Tbx3*. *Tbx2* is the likely candidate, being the only *Tbx*-family member whose expression is highly correlated with onset of the eye field.

**Fig. 2.**
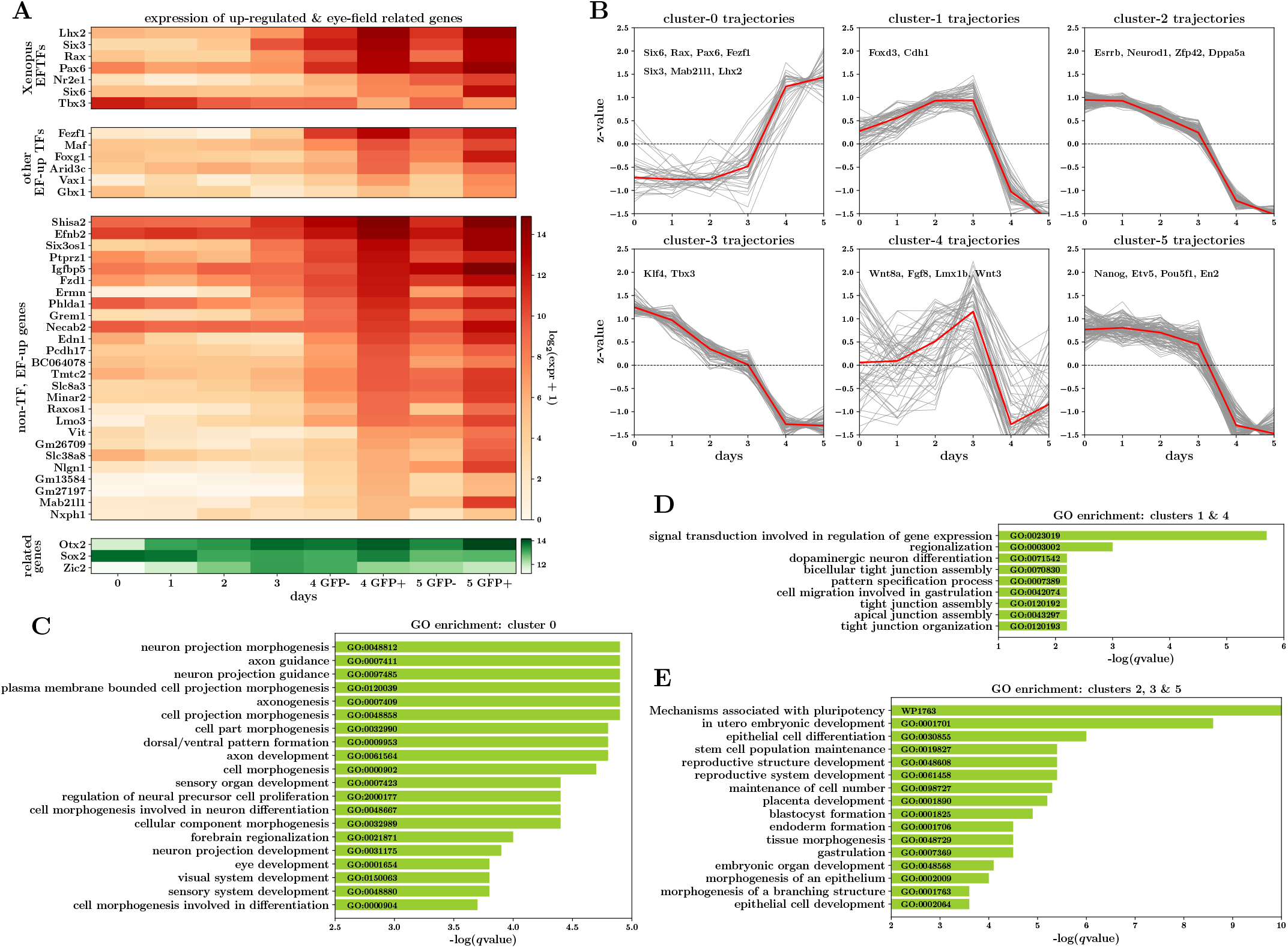
RNA-seq analysis reveals coordinated up-regulation of canonical EFTFs during optic vesicle development. **A.** Expression heatmap displaying gene-expression changes across the OV timecourse, for canonical eye field genes, differentially up-regulated (EF-up) genes and genes known to have an important role in early eye development. **B.** Clustering of EF-up and EF-down gene trajectories reveals seven clusters with three underlying overall patterns across the timecourse. **C–E.** Gene ontology enrichment analysis of sets of genes within the three trajectory patterns revealed by clustering.

In parallel to the up-regulation of this extended set of putative eye field genes, the cell-state transition is also characterised by the *down*-regulation of a larger separate set of “EF-down” genes (448) across the day3-to-day4/day5 transition (Fig.S2A). Many of these genes are known major players in pluripotency maintenance, including *Nanog, Pou5f1* and *Dppa5a*, and although they are down-regulated across the state-transition, they are also consistently down-regulated across the full timecourse of organoid differentiation. This suggests that successful specification of the eye field requires both the turning-on of the network of eye field genes as well as the switch-off of genes controlling pluripotency. It is noteworthy that *Sox2* and *Otx2*, known to be crucial in eye development are not differentially expressed across this critical time-point (Fig.2A).

We next further probed our RNA-seq data to see whether any of the EF-up and EF-down genes displayed significant changes between earlier time-points. Performing a similar differential expression analyses, we find that 9 (out of 165) genes up-regulated between day 2 and day 3 samples overlap with the EF-up genes, including the TFs Rax, *Lhx2, Six3* and *Fezf1*. The fact that these key factors are significantly up-regulated so early in OV development (even prior to distinct GFP signal), may point to these TFs acting as *very early* eye field determining factors. Interestingly, 12 genes up-regulated between day 2 and day 3 actually overlap with EF-down genes, including *Wnt3, Wnt8a, Fgf8* and *Lmx1b*, pointing towards the eye field cell-state requiring a repression of gene-expression programs turning on earlier in the differentiation cascade. Indeed, this has close parallels to what is thought to happen *in vivo*, where the EFTFs, as well as promoting ocular fate, are also involved in repressing gene-expression programs characterising other neural cell-states, such as di-encephalic, mid-and hind-brain states, early in development (Roy, 2013; Fish et al.,2014; Liu et al., 2010; Takata et al., 2017).

Finally, to explore whether there are general patterns to the trajectories of differentially expressed genes, we performed a clustering analysis on the trajectories of the EF-up and EF-down genes, and of genes differentially expressed between day 2 and day 3 (Fig.2B). This analysis resulted in 6 clusters which roughly fall into three trajectory patterns: genes in clusters 0 are steeply up-regulated after day 3 (all the EF-up genes fall into this category), clusters 2, 3 and 5 contain genes with expression gradually down-regulated with time, and clusters 1 and 4 which contain genes with more transient expression, being broadly up-regulated until day 3 and then sharply repressed after day 3. Whilst the focus of our work is to gain a better understanding of the mechanisms underpinning the gene trajectories in cluster 0, this analysis provides further evidence that the stable onset of the EF identity requires both the up-regulation of a set of genes as well as suppression of other gene-expression programs which would otherwise drive cells to alternative fates. A gene ontology (GO) enrichment analysis (Zhou et al., 2019) on the sets of genes displaying these three broad patterns provides an additional layer of evidence that the stable onset of the eye field is characterised by a balance of very different geneexpression programs: the up-regulation of a gene expression program specifically related to eye and neuron development (Fig.2C), and in parallel the down-regulation of gene-expression programs characterising pluripotency, non-neuroectodermal differentiation (Fig.2E) as well as possibly cell signalling and migration (Fig.2D). In summary, these first analyses of RNA-seq data generated from the timecourse of optic vesicle organoid development, show that this model system is not only a robust and relevant model system with which to study mammalian eye field specification, but also one that can shed light on how other cell-fates during early brain development are suppressed.

### Dynamic ATAC-seq peaks local to eye field genes suggest enhancer-driven logic

Changes in the expression of genes characterizing cell-state transitions, in development as well as in disease, are thought to be largely controlled by changes in DNA-accessiblity and binding of TFs at *cis*-regulatory elements genomically local to those genes. Therefore, to gain insight into the regulation of the transcriptional changes identified as important for the eye field transition, and specifically to identify candidate TF-*cis*-regulatory element pairs that control the expression of EF-genes, we performed bulk ATAC-seq (Buenrostro et al., 2013) experiments on days 0, 1, 2 and 3, as well as GFP positive and GFP negative cells on day 5. After quality control and peak calling, we identified a consensus set of 361,867 genomic regions displaying a significant accessibility peak on one or more days. Mirroring the strategy used to identify interesting genes in the RNA-seq analysis, we look for peak regions that are dynamic (absolute log_2_ fold-change > 1.5) across day 3 and day 5, finding 7,782 and 53 such peaks with increased and decreased ATAC-seq signal respectively. After excluding peaks overlapping exons (TFs typically bind non-exonic regions), and annotating peaks as promoter or non-promoter (intergenic or intronic), we find that this set of dynamic peaks is comprised largely (~ 97%) of non-promoter peaks. This feature of our ATAC-seq data is consistent with previous work that indicates changes in the chromatin accessibility landscape during early development predominantly occur in non-promoter regions (Reddington et al., 2020), suggesting an enhancer-driven logic to cell-state changes. Moreover, computationally linking these dynamic ATAC-seq peaks to genes in an unbiased manner, using a GREAT analysis (McLean et al.,2010)), indicates gene-peak associations for 22 of the 37 up-regulated and 161 of the 448 down-regulated genes. This suggests that the observed changes in chromatin accessibility may be functionally relevant in the regulation of differentially expressed genes characterising the eye field.

Although genome-wide chromatin-accessibility dynamics is of interest in itself, our key goal is to uncover changes in chromatin accessibility that play a direct role in regulating the changes in gene-expression identified using the RNA-seq data. To identify *cis*-regulatory elements relevant to the putative eye field genes, we focus our attention on peak regions that lie within the topologically-associated domains (TADs) (Dixon et al., 2012;Nora et al., 2012) containing those genes (using TADs defined using mESCs Hi-C data (Bonev et al., 2017)). This approach is motivated by the observation that TADs appear to be remarkably stable across development and tissue-types and that gene regulatory elements generally lie within the TAD containing the target gene (Symmons et al., 2016). Upon narrowing down to this set of peaks, we find that 98% and 93% of dynamic peaks in up- and down-regulated eye field TADs respectively, correspond to non-promoter peaks. By contrast, peaks overlapping promoters of many eye field genes, including *Rax*, *Pax6* and *Lhx2*, are accessible from day 0, despite very low or zero corresponding gene expression (Fig.3B and Fig.S3A). Indeed we find that while promoter regions of the putative EF genes display higher median ATAC-seq signal than corresponding dynamic non-promoters (Fig.3A, *p* < 10^−229^, Mann-Whitney U-test), the latter have distinctly more variable trajectories across the timecourse of our experiment (Fig.3A, p < 10^−243^, Mann-Whitney U-test, and Fig.3B). The above observations are again suggestive of an overall picture whereby gene-expression changes underlying the transition to an ocular-fate are orchestrated to a large extent via changes in the accessibility and TF-binding at distal and intronic regulatory elements.

**Fig. 3.**
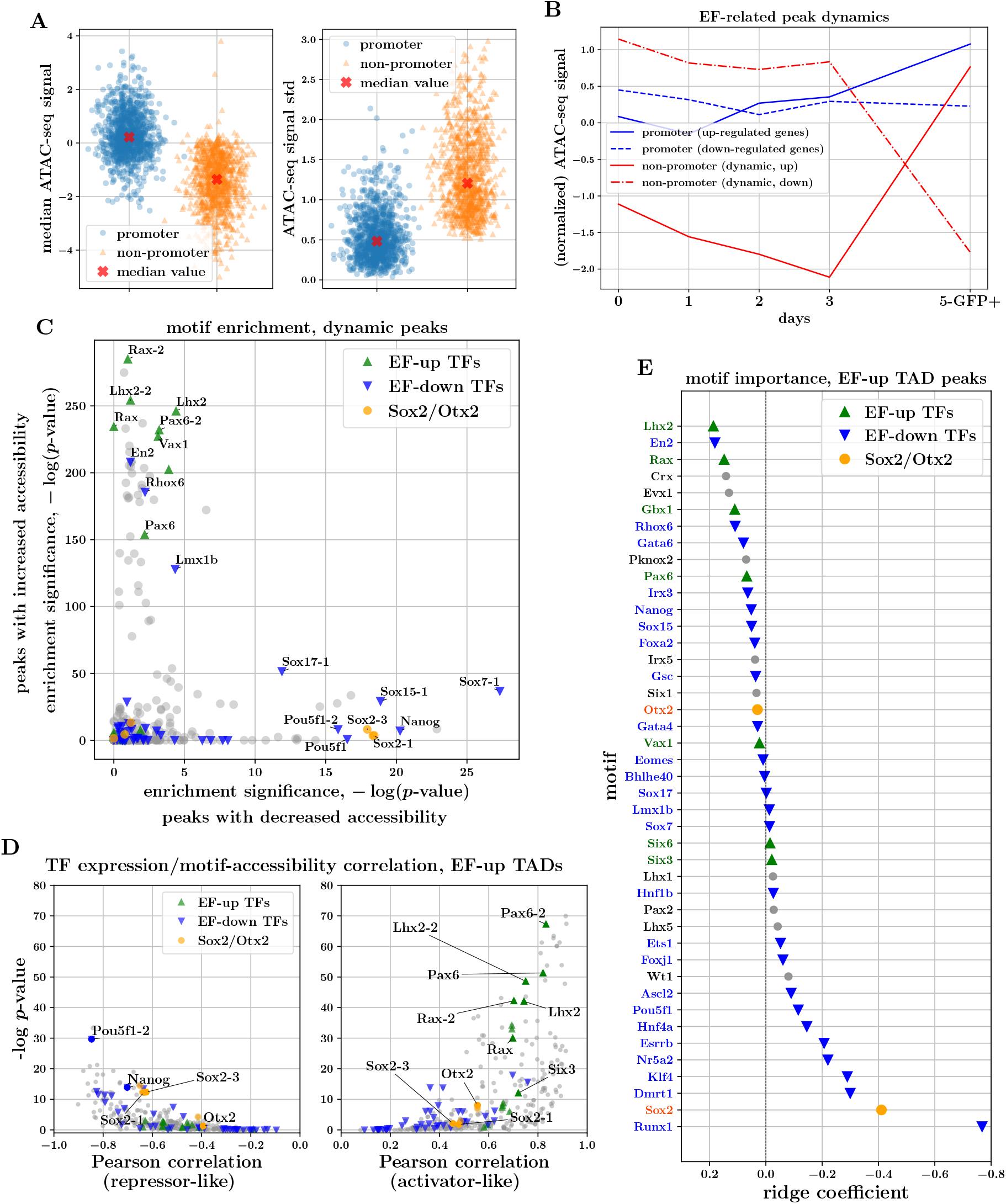
Characteristics and sequence analysis of ATAC-seq peaks associated with the transition to eye field state point to enhancer-driven logic and a key regulatory role for canonical EFTFs. **A.** Mean and variability of ATAC-seq signal in promoter and non-promoter peaks associated with eye field genes. **B.** ATAC-seq signal trajectories of up- and down-regulated gene promoters, as well as of dynamic non-promoter peaks within EF TADs. **C.** TF-motif enrichment analysis of peaks with increasing (y-axis) and decreasing (x-axis) accessibility signal across day3-day5 transition. **D.** TF-expression/TF-motif-accessibility correlations for peaks within EF-up TADs. RHS and LHS plots illustrate the median across positive and negative correlations respectively, indicative of activator-like and repressor-like behaviour of the respective TFs. **E.** Coefficients of logistic-regression model trained to predict opening-vs-closing behaviour of ATAC-seq peaks within EF-up TADs, using presence of TF-motifs as input covariates. Magnitude of coefficients indicative of importance of respective TF-motifs for these predictions.

### Motifs of up-regulated eye field genes are enriched in and predictive of regions of dynamic chromatin

As TFs bind DNA in a sequence-biased manner, analysing the sequence content of accessible genomic regions provides preliminary evidence of the classes of TFs that may be involved in gene regulation. First, we performed a motif enrichment analysis on the set of peaks identified as dynamic across the day3-day5 transition (scanning peak regions with motif-PWMs from the JASPAR2020 database (Fornes et al., 2020) using `fimo` (Grant et al., 2011)). We find that peaks with increasing accessibility have a strong enrichment for the motifs of the canonical EFTFs *Rax*, *Lhx2* and *Pax6*, whilst peaks with decreasing accessibility appear enriched for motifs of pluripotency factors such as *Pou5f1* and *Nanog* (Fig.3C). This provides the first clues that the TFs characterising the transcriptomic state-transition may also play an important role in re-shaping and/or activating regulatory regions of chromatin. Furthermore, to assess whether the evidence from the motifenrichment analysis of genome-wide dynamic peaks is consistent with accessibility dynamics in peaks related to the EF-up genes, we performed a multivariate determination of motif-importance for peaks within EF-up TADs. Specifically, we fit a logistic-regression model to predict whether the accessibility signal of a peak increases or decreases across the day3-to-day5 transition, using as input the binary motif presence of differentially expressed TFs. We find that the motifs corresponding to up-regulated TFs, and in particular *Lhx2* and *Rax*, are most predictive of opening chromatin, whilst motifs corresponding to pluripotency factors are predictive of closing chromatin (Fig.3E). The overall picture of key EFTFs and TFs associated with pluripotency being predictive of opening and closing chromatin peaks respectively, appears to be a general one genome-wide as we find very similar coefficient patterns when we repeat the modelling exercise for genome-wide dynamic peaks instead of peaks in EF-up TADs (Fig.S3B).

Interestingly, this multivariate approach also reveals that there are a number of down-regulated TFs, such as *En2*, *Gata6* and *Nanog* that are predictive of increases in accessibility. This may well be related to the fact that motifs of many TFs are similar and have overlapping occurrences genome-wide. This common issue with standard motif analyses makes it particularly difficult to gain deeper insights into the molecular logic of interest. One approach to move closer towards a more reliable set of regulatory TFs is to use ATAC-seq together with RNA-seq data. Working with the assumption that a TF that alters the accessibility of regulatory elements must be expressed to do so, there ought to be a significant correlation between the accessibility of its DNA binding motif and its gene expression. Using an approach similar to Corces et al. 2016 and Berest et al. 2019, for each TF we compute the correlation between its expression and the accessibility of each peak containing the TF-motif, and then compare the distributions of positive and negative correlations against relevant background distributions to determine significance. We find that for non-promoter peaks within EF-up gene TADs, the canonical EF genes *Rax*, *Pax6*, *Lhx2* and *Six3* as well as *Otx2* display a significant positive expression-accessibility correlation (Fig 3D, median corr > 0.5, p < 10^−15^ Wilcoxon rank-sum test), suggesting activator roles for these transcription factors. The same analysis reveals potential repressive roles for the pluripotency factors *Pou5f1, Nanog* and *Sox2*, which display significant negative expression-accessibility correlation (Fig 3D, median corr < −0.5, p < 10^−15^ Wilcoxon rank-sum test) for peaks within EF-up gene TADs. This is in line with evidence suggesting that as well as being important for activating CREs maintaining pluripotency, these genes also act in a repressive manner to block activation of elements crucial for the up-regulation of differentiation genes (Thomson et al., 2011; Wang et al., 2012). Applying this analysis to peaks in EF-down TADs (Fig.S3C), we find, as expected, that pluripotency factors there appear to have activator-roles, in line with knowledge that these TFs bind enhancers that up-regulate their expression. Surprisingly, we also find that peaks within EF-down TADs containing EFTF motifs tend to have have accessibility signal that correlates well with EFTF expression (Fig.S3C). This observation provides a hint that, alongside being crucial for specifying the eye field geneexpression program itself, the EFTFs may also play a role at CREs repressing genes characteristic of alternative gene-expression programs (consistent with evidence that EFTFs repress alternative cell fates *in vivo* (Roy, 2013;Fish et al., 2014;Liu et al., 2010; Takata et al., 2017)).

Taken together, the above TF-motif guided analyses and modelling of chromatin-accessibility changes consistently point to the hypothesis that the stable up-regulation of EF-genes is enabled through the gain of binding of a small set of EFTFs at nonpromoter CREs that are becoming increasingly accessible, in genomic domains that are local to those genes. Additionally, our analyses provide evidence that TFs down-regulated across OV development, act in a repressive manner, and thus the loss of their corresponding binding at elements with decreasing accessibility, perhaps through a loss of ‘silencing’, may also be contributing to the activation of the EF-up genes.

### Footprinting analysis provides evidence of differential binding of key TFs

Detectable ‘footprints’ or relative depletions in chromatin accessibility signal at regulatory elements, can result from DNA being protected from transposase cleavage due to the presence of bound TFs, and can therefore provide further evidence of binding of a given TF. Footprinting analyses also have the potential to distinguish more subtle effects, such as changes in the distribution of TF-binding across conditions, which can be particularly useful if regulatory elements do not display overall changes in accessibility. We applied the TOBIAS framework (Bentsen et al., 2020) to our ATAC-seq data to identify putative changes in binding and foot-printing of expressed TFs, both in genome-wide peaks as well as in peaks within EF-TADs. To do this `TOBIAS` computes a ‘footprint score’ for each occurrence of a TF-motif (within the peaks of interest). This represents the combined effect of how well the underlying occurrence sequence matches the TF-motif together with the evidence of protein binding from the ATAC-seq data (relative depletion in chromatin accessibility at the motif occurrence).

We first performed a differential binding analysis between day 3 and day 5 on peaks genome-wide, which compares footprint scores for every occurrence of a TF-motif between days (providing an aggregate pattern across all occurrences of a motif within peaks of interest). This reveals that some of the largest increases in motif-binding correspond to EF-up TFs, most notably Rax, Pax6 and Lhx2 motifs (Fig.4A), and similarly, that some of the most significant decreases in motif-binding correspond to EF-down TFs, such as Pou5f1 and Dmrt1. Consistent with our motif analysis in the previous section, Six3 does not appear to have a strong increase in binding across the day3-day5 transition, perhaps indicating that Six3 plays a regulatory role in eye field specification that is different to direct binding to regulatory elements of EF genes (e.g. Six3-mediated Wnt8 suppression has been shown to be important for normal neuroretina development (Liu et al., 2010)). Interestingly, we find that there is evidence of increased binding of Otx2, consistent with the hypothesis that, together with Sox2, Otx2 is required for the coordinated up-regulation of the EFTFs and in particular leads to activation of *Rax* expression (Danno et al.,2008).

**Fig. 4.**
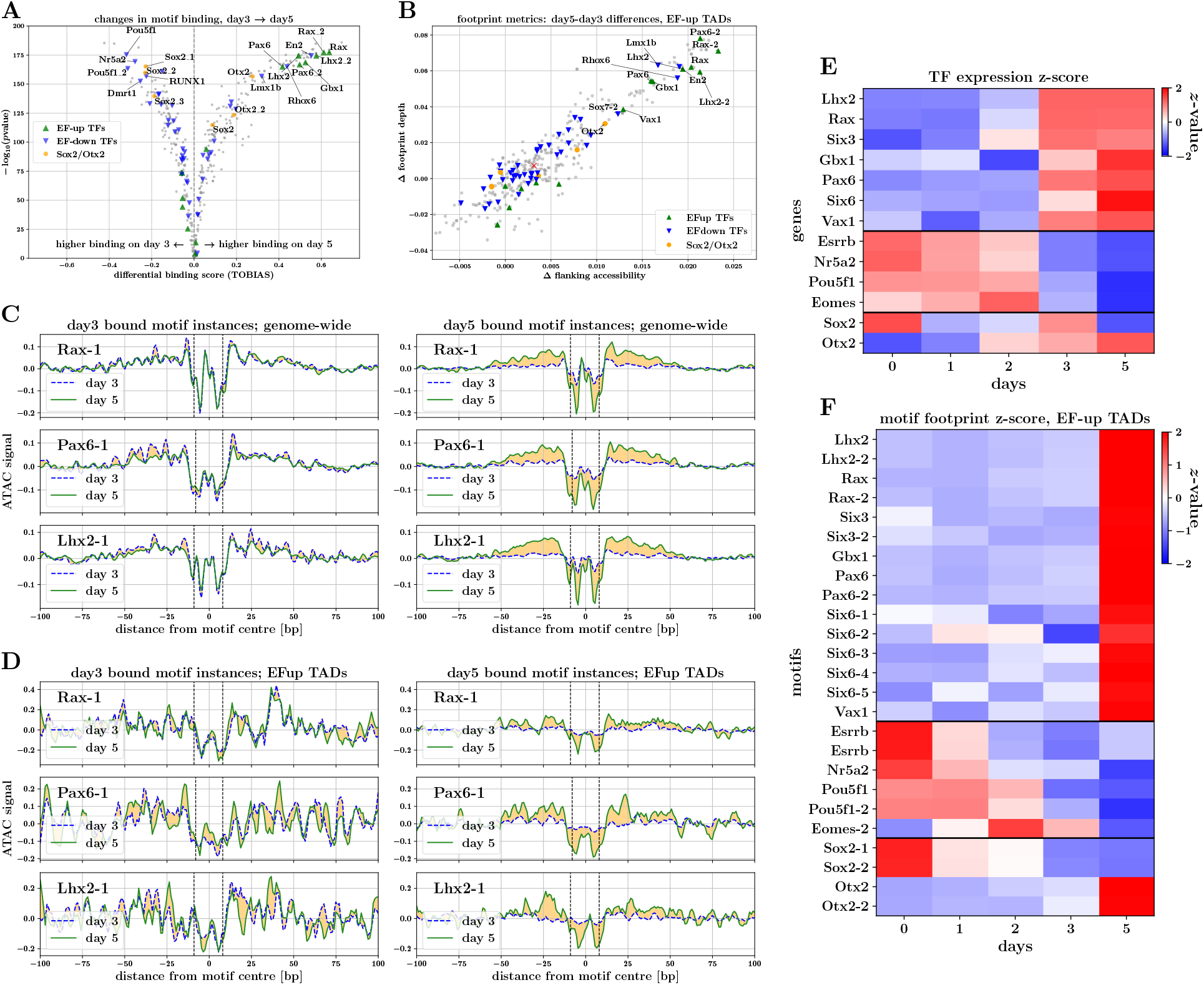
Footprinting analysis provides evidence of direct interactions of EFTFs with DNA. **A.** Day3-day5 differential binding scores for motifs of expressed TFs. **B.** Scatterplot of day3-day5 differences in TF-motif footprint depth versus flanking accessibility, for peaks in EF-up TADs. **C. & D.** Aggregate footprint signals for *Rax, Pax6* and *Lhx2* motifs. Plots illustrate aggregate signal on day 3 and day 5 for motif-instances predicted to be bound on day 3 (left) and day 5 (right). **C.** Aggregate footprint signals for genome-wide motif instances. **D.** Aggregate footprint signals for EF-up TAD motif instances. **E. & F.** Expression and motif footprint score trajectories for TFs displaying strongest expression-footprint correlations in EF-up TADs. **E.** Heatmap of TF-expression values *z*-transformed across timecourse. **F.** Heatmap of TF-motif footprint scores in EF-up TAD peaks, *z*-transformed across timecourse (median across peaks).

Furthermore, considering footprint scores across the OV-development timecourse, we assume that a TF playing a regulatory role via direct binding to DNA, should display footprint scores that correlate well with its expression. To investigate which motifs show such correlated footprint-score/expression behaviour, for each motif we compute the correlation of the footprint score of each occurrence with the corresponding TFs expression. The set of motifs that display a median correlation > 0.5 in EF-up-TAD peaks contains all key EF-up TFs (visualized in Fig.4E using z-transformed expression or median footprint-score), as well as Sox2 and Otx2. When comparing the footprint score dynamics in EF-up-TAD peaks and genome-wide, we find that there is generally good agreement overall for most motifs. Notably, however, footprint scores for Six3 and Six6 motifs break this trend and don’t correlate well with their corresponding expression when looking at occurrences genome-wide (Fig.S4F).

Aggregating ATAC-seq signal around multiple occurrences of a given motif, allows for the visualization and quantification of TF-footprints. To unpick large changes in footprinting between day 3 and day 5 in an unbiased manner, we computed the depth and flanking accessibility for each motif, for genome-wide and EF-up TAD peak occurrences, on each day. Comparing the changes in both footprint depth and flanking accessibility between these time points, we find that in EF-up peaks (Fig.4B) as well as genomewide (Fig.S4B), motifs of the up-regulated TFs Rax, Pax6 and Lhx2 (as well as of Vax1 and Gbx1) display some of the largest positive changes amongst all motifs of expressed TFs. Curiously, similarly large increases in these scores are also found for the down-regulated TFs Lmx1b, En2 and Rhox6 (drawing parallels to what we found using our approach to model opening/closing chromatin peaks using motifs, Fig.3E).

Finally, to confirm that the above large increases in footprint depth scores do correspond to visually discernible footprints, we visualize the aggregate ATAC-seq signal at and around occurrences of a given motif. In order to pull apart changes between days, we find it instructive to compare the aggregate signal for motif occurrences that are predicted by `TOBIAS` to be ‘bound’ on day 3 and on day 5. Whilst there appears to be little difference in aggregate signal between days, for occurrences predicted to be bound on day 3, we find a striking increase in footprint depths for Rax, Pax6 and Lhx2 motif occurrences predicted to be bound on day 5. This holds both for peaks genome-wide as well as in occurrences within peaks in EF-up gene TADs (Fig.4C,D). This provides further evidence that these TFs appear to be regulating the onset of eye field specification by binding elements in the genomic locality of EF-up genes. We also find that aggregate footprints for Pou5f1, Sox2 and Otx2 show differences (albeit less striking) that are consistent with the picture emerging that establishment of the eye field appears to be correlated with a loss of binding of pluripotency factors and a gain of binding of Otx2 (Fig.S4C,D).

### Peak and TF specific footprint-score changes identify candidate *cis*-regulatory elements

The motif and footprint analyses described above provide consistent evidence that the canonical EFTFs *Rax*, *Pax6* and *Lhx2* likely play a direct regulatory role in establishing a stable eye field, specifically through binding non-promoter elements in accessible regions within TADs containing EF-up genes. As a final step to characterising the regulation of EF-up genes, we wish to move from this broad picture of regulation towards identifying candidate gene-specific regulatory elements through which these key TFs act. For a given EF-up gene, we have approached this by searching for ATAC-seq peak regions within the TAD containing the gene, which display large changes in footprint scores for motifs of any of these three key TFs, as well as of *Sox2* and *Otx2*. In particular we look for changes between days 1, 3 and 5 that either mirror the up-regulation of genes or which may point to a loss of potential repression.

This approach is visualized in Fig.5 for peaks within the *Rax* (Fig5.A&B) and *Six6* (Fig5.C&D) TADs, where the difference in footprint scores for each motif across the day3-day5 and day1-day3 time-points are illustrated in the heatmap. Focusing on changes between day 3 and day 5 within the *Rax* TAD (Fig.5A), the heatmap reveals highly suggestive footprint score changes across the majority of the key TFs, in three peak regions (peak ids 165713, 165714 and 165716), which mirror the up-regulation of the *Rax* gene itself across these time-points (Fig.S5A). A similar pattern is seen within the *Six6* TAD (Fig.5C) where there are two peak regions (peak ids 73704 and 73706) showing high footprint score changes and increasing accessibility (Fig.S5B). No striking increases in footprint scores are identified across the day1-day3 transition, however there are some peaks that show a decrease in scores particularly for the Sox2-related motifs (most notably peak id 165695 proximal to *Rax*). We point out that many of these changes in footprint score are correlated with the log_2_-fold-changes in overall ATAC-seq signal for each peak. We find similar observed patterns of signal, highlighting a small number of peaks as candidate regulators for each EF-up gene (changes in *Pax6*, *Lhx2* and *Six3* TADs are shown in Fig.S6).

**Fig. 5.**
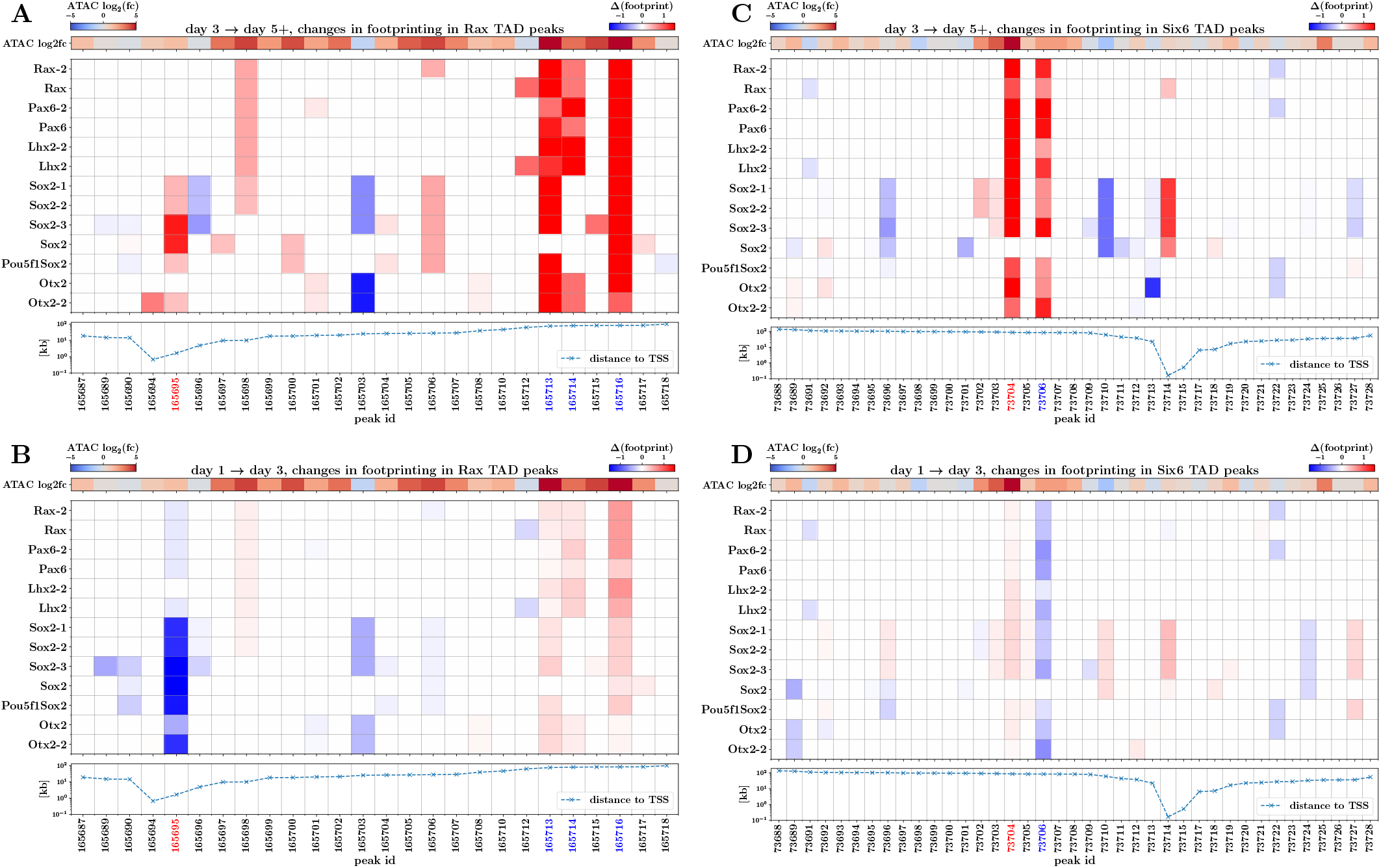
Changes in motif footprint scores of key TFs identify candidate *Rax* and *Six6 cis*-regulatory elements. i **A.** Heatmap of day3-to-day5 changes in footprint scores for motifs of key TFs at peaks contained in Rax TAD region. Also illustrated are associated changes in normalized ATAC-seq signal for each peak (upper panel), and distance of each peak from the Rax TSS (lower panel). **B.** Same as **A.** but displaying changes between day1-to-day3 changes. **C. & D.** are same as **A.** and **B.** but displaying changes in peaks contained in *Six6* TAD region. Peak-IDs highlighted in red indicate those chosen for CRISPR-perturbation whilst peak-IDs highlighted in blue indicate other putative cis-regulatory elements with interesting footprint score changes (see also Fig.S5).

We have selected two candidate regulatory elements, one for *Rax* and one for *Six6* (highlighted as red-coloured peak ids in Fig.5) to test for enhancer-like behaviour for the genes they are linked to. The candidate peak proximal to *Rax* is accessible from Day 1 (Fig.6A&S5A), and although its accessibility remains constant over time, the footprint score changes indicate a loss of Sox2 binding from day 1 to 3 and then a gain of binding from day 3 to 5 suggesting there could be a switch from repressive-to-activating function of this peak region. Additionally, this peak overlaps with a putative regulatory element identified by Dannoet al., 2008 in *Xenopus* (discussed further in the next section). The candidate peak selected for *Six6* shows a simpler pattern of accessibility being closed at day 1 and becoming accessible by day 5 (Fig.6D&S5B), and there is a high correlation between *Six6* gene expression and peak accessibility.

**Fig. 6.**
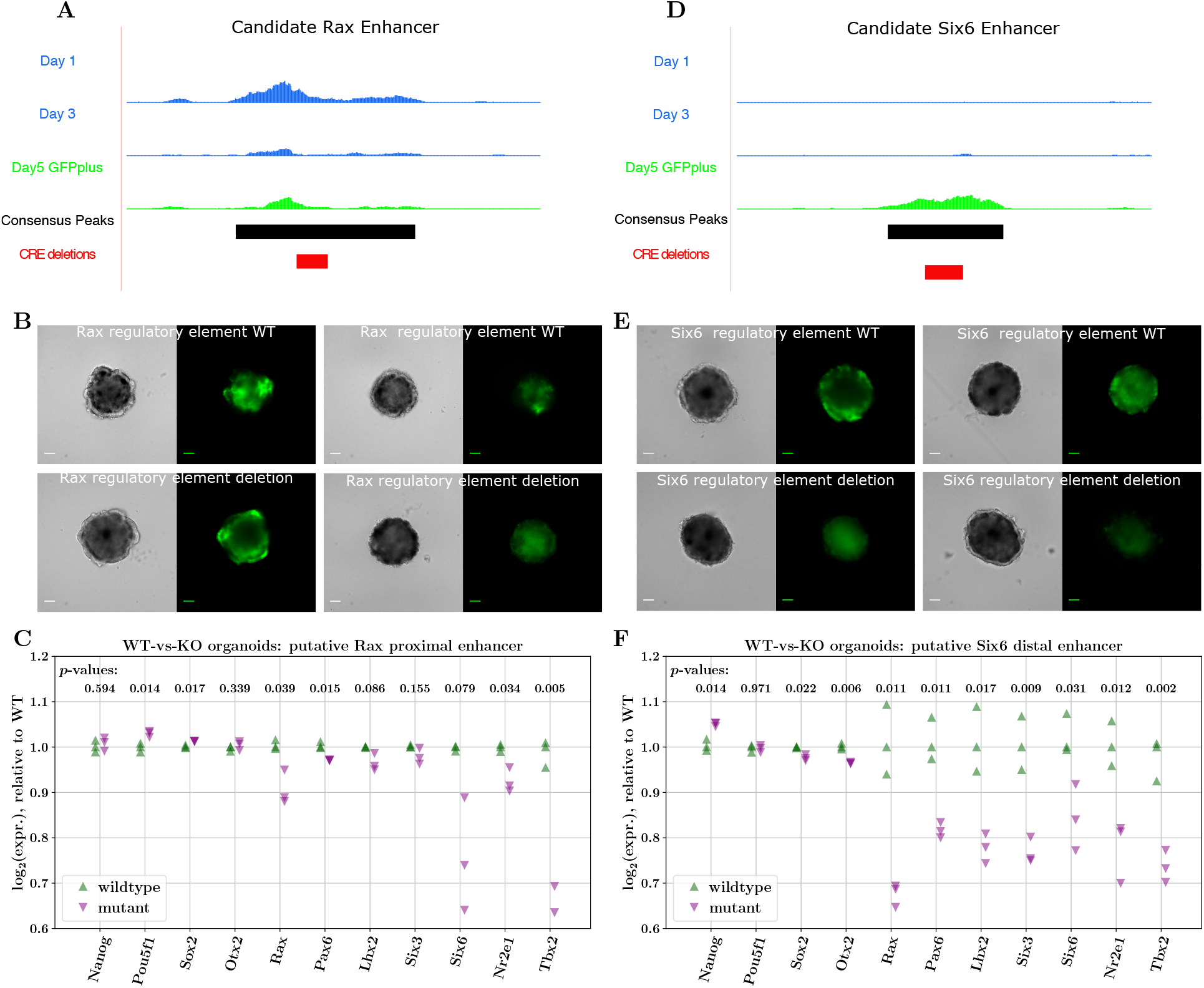
Changes in expression of key TFs upon disruption of putative enhancers of EFTFs. **A. & D.** Bigwig tracks of candidate *Rax* (proximal) and *Six6* (distal) enhancers, displaying changes in accessibility signal across days 1, 3 & 5, the consensus peak regions and the regions knocked-out by CRISPR. **B. & E.** Two organoid images derived from wildtype (top panel) and mutant CRISPR cell-lines representing the variability seen in structure and GFP expression of organoids from these differentiations. Scale bars: 100 *μ*m. **C. & F.** NanoString quantification of gene expression differences between wildtype and CRISPR– mutant organoids. Subfigures **C.** and **F.** illustrate differences upon perturbation of putative proximal Rax enhancer and distal *Six6* enhancers respectively. Statistical testing of differences in log_2_ expression between wildtype and CRISPR-mutant organoids performed using Welch’s t-test.

We used CRISPR-Cas9 technology to generate stable cell lines with deletions encompassing the binding motifs of key TFs that were identified in the candidate regulatory elements. In the candidate *Rax* regulatory peak we introduced a 113bp homozygous deletion that disrupts two Sox2 motifs and a Sox2-Pou5f1 motif as well as an Otx2 and a Pax6 motif. This includes the Sox2 and Otx2 motifs identified as regulating *Rax* expression in *Xenopus* Danno et al., 2008. For the *Six6* peak a 140bp homozygous deletion disrupted one site that is predicted to be bound by Lhx2, Rax and Pax6, as well as two Sox2 binding motifs and a second Lhx2 motif. We then used these mutant cell lines to generate OV organoids (Fig.6B&E). The organoids grown from these cells were considerably more variable than previous batches of organoids (Fig.S7&S8), meaning no clear conclusions could be drawn from looking only at organoid structure and GFP expression. To test whether perturbation of these putative gene-specific cis-regulatory elements had an effect on the expression of the expected target gene, we performed digital gene expression (NanoString) analysis (Geiss et al., 2008) in triplicate on organoids at day 5 of differentiation, grown from cell lines that had no detected mutations introduced by the CRISPR machinery (referred to as wildtype) and cell lines with the mutations described.

Comparing the expression of key genes in wildtype and mutant organoids, we find that disrupting the candidate elements for both *Rax* and *Six6,* results in the down-regulation of the expected target gene (Fig.6A, −11% for *Rax* and −16% for *Six6* log_2_-expression, respectively). Notably, for both mutants, *Tbx2* is one of the most down-regulated genes adding further support to our hypothesis that *Tbx2* is the mammalian equivalent of *Tbx3* in *Xenopus*. Furthermore, we also find that in both *Rax* and *Six6* putative enhancer mutant organoids, the expression of *all* key EFTFs is reduced, as might be expected if this set of key TFs really do co-regulate each other (directly or indirectly) within a GRN.

### Regulatory switches responding to ratios of TF concentration may trigger initial EFTF activation

An important question that is difficult to disentangle directly with the resolution of our experiments and data is via what mechanism is the expression of key EFTFs actually triggered? To answer this question, we note that experiments (albeit in model systems different to the mESC-derived organoids) in which *one* of the TFs *Rax*, *Pax6*, *Lhx2* and *Six3* has been knocked-out or rendered null, have shown that expression of the non-perturbed genes is actually activated in the appropriate eye field regions (Zhang et al.,2000;Fish et al., 2014;Grindley et al., 1995;Bernier et al., 2001;Yun et al., 2009; Roy, 2013; Liu et al., 2010). Maintenance of this expression is observed to be significantly de-stabilised. This provides evidence towards the hypothesis that the molecular events leading to the first onset of individual EFTF expression are not dependent on the EFTFs themselves.

The observations in the experiments discussed above, would mean that the molecular events leading to the switch-on of *Rax*, and specifically, the required changes at regulatory elements that trigger *Rax* expression, do not necessarily depend on or involve the other EFTFs (*Pax6*, *Lhx2*, *Six3*). Related to this, the work of (Danno et al., 2008), showed in *Xenopus* animal cap cells, that Sox2 and Otx2 proteins interact to activate *Rax* expression, by cobinding a conserved 35bp region within a cis-regulatory element ~ 2kb upstream of the *Rax* promoter. The authors also propose that activation of this regulatory element is sensitive to the ratio of Sox2 and Otx2 concentrations (Fig.S9B).

As mentioned in the previous section, the OV organoid ATAC-seq data we have generated in this study contains a peak that overlaps with the conserved element identified by Danno et al., 2008. Intriguingly, we find that this putative regulatory element is accessible (with no large changes in overall accessibility) throughout the organoid development timecourse, and thus appears to be primed for activation. Furthermore, if it is the ratio of Sox2 and Otx2 that leads to the activation of this *cis*-element, then we would expect to see sizeable differences in Sox2 and Otx2 binding between OV development days. Indeed this is what we do find for Sox2, where the footprint score for Sox2 motifs decreases from day 1 to day 3 and increases again from day 3 to day 5, specifically at this regulatory element, possibly indicating changing of Sox2 binding partners across the timecourse (Fig.5A&B). The predicted binding ratios for Otx2 at this element are less pronounced, although they do increase between day 3 and day 5, in line with the working hypothesis. Finally, we report that by exploring two publicly-available Sox2 and Pou5f1 ChIP-seq datasets (Xiong et al., 2022; Avsec et al., 2021) generated from E14 mESC-cells (roughly equivalent to day 0 organoids in the OV timecourse), we find peaks of Pou5f1 signal at this cis-element (Fig.S9A). This suggests a mechanism whereby, although this element is accessible early in OV development, it is repressed by Pou5f1 binding and thus not able to activate *Rax* expression. As the expression of *Pou5f1* is down-regulated and the expression of *Otx2* increases, Sox2 may exchange binding partners (from Pou5f1 to Otx2) and in so doing, activate this putative enhancer and consequently also the expression of *Rax*.

Finally, we note that should the above hypothesis of regulatory switches be a model that generalizes beyond just the activation of *Rax*, i.e. that *cis*-regulatory elements acting as triggers of EFTF expression are ubiquitously accessible and rapidly change behaviour from silenced/repressive to activating by responding to differences in concentrations of different TFs, then these would generally be challenging to identify through a standard strategies of looking for correlated changes in overall accessibility.

## DISCUSSION

By using an optic vesicle organoid model to enrich for cells transitioning to an optic fate, we have generated bulk transcriptomic and chromatin accessiblity timecourse data from a specific and very early developmental event critical for mammalian eye development. Joint computational analyses of these data have revealed three key messages regarding the molecular events controlling eye field specification. Firstly, the transition to the eye field state in mESC-derived OV organoids is characterised by the up-regulation of a small set of genes, including a core set of transcription factors known to be necessary for eye development through studies in *Xenopus*. Secondly, the patterns extracted from our RNA-seq data indicate that when this state-transition occurs, not only is this core set of EFTFs up-regulated, but in parallel, other geneexpression programs important in early development, such as those controlling cellular pluripotency and cellular signalling are down-regulated or carefully balanced. Thirdly, analysis of the dynamics and sequence composition of chromatin accessibility peaks strongly suggests that the key EFTFs – particularly *Rax*, *Pax6* and *Lhx2* – act to control the up-regulation of the EF genes through binding to (primarily) non-promoter regulatory elements in the vicinity (TADs) of those genes.

By performing footprinting analyses for key TFs in TADs of important genes, we have identified putative gene-specific *cis*-regulatory elements that appear important for regulating the gene-expression changes characterising the onset of the eye field. Furthermore, by perturbing a subset of these candidate regulatory elements via CRISPR and generating mutant cell lines, we have started to validate the regulatory potential of these elements in controlling EFTF expression in OV-organoids. Our work demonstrates the power and practicality of using such an organoid system to generate and test specific hypotheses regarding the regulation of cell-state transitions relevant to critical events in mammalian development.

Although our data does not have the resolution to definitively identify what may trigger the expression of individual EFTFs, we have presented evidence to suggest that this may arise, at least in the case of *Rax*, from the replacement of repressive factors (such as Pou5f1) with activating factors (such as Otx2) at regulatory elements that are ubiquitously open across the timecourse. We have also outlined that disentangling the mechanisms behind geneexpression or cellular-state changes that are triggered through the action of TFs at such regulatory elements, can indeed be very challenging to understand through standard analyses correlating peak-accessibility with gene-expression. However, footprinting analyses do have the potential to provide richer insights into these more subtle mechanisms of regulation.

In undertaking the work presented here we have encountered a number of intriguing open questions regarding eye field specification which require further study to resolve. In particular, it is pertinent to ask whether the eye field directly stems, in a sequential manner, from a group of well-defined progenitor cells within the neural plate, or whether individual neuroectodermal cells spontaneously acquire ocular fate and are only then organised into a coherent eye field through migration and cell-signalling events? Furthermore, are these processes similar *in vivo* and in the organoid system? To further probe the emergence of the eye field, it would be fascinating to investigate the extent to which the EFTFs act in a cell-autonomous manner, regulating the genes of the cells in which they are endogenously produced, or whether there are non-cell-autonomous components to gene regulation (paracrine signalling). Recent rapid technological progress in single-cell technologies – both multi-omics and imaging – will enable exciting future research in these directions. Finally, in order to bring this research closer to having translational potential, similar analyses must be performed using organoids derived from human ESCs – indeed such a line of work has the potential of identifying and testing mechanisms crucial to early human eye development and disease that are currently impossible to study.

## MATERIALS AND METHODS

### ES Cell Maintenance and Organoid Culture

Rx-GFP mouse ES cells were maintained and SFEBq differentiation of OV organoids was performed as described previously (Wataya et al., 2008; Eiraku et al., 2011). Briefly, mESCs were dissociated and 4500 cells were plated in differentiation media (GMEM supplemented with 1.5% KOSR (KSR media) or a growth factor-free chemically defined medium (CDM media) for KO organoids and initial organoids respectively) in a low cell adhesion 96 well plate. This is defined as day 0. On day 1, growth factor reduced Matrigel was added to a final concentration of 2%. Differentiation status was assessed using fluorescent imaging to detect the presence of GFP as a measure of *Rax* expression.

### RNA-Seq Assays

On days 0–5 of optic vesicle differentiation, triplicate samples of cells were collected from 24 organoids which were dissociated and sorted using FACS to select for live single cells. Samples at day 4 and 5 were also sorted on the basis of GFP expression giving a GFP positive and negative sample for each of these time points. RNA was isolated using the Zymo Direct-zol RNA MicroPrep kit following manufacturer’s instructions. Quality and integrity of the RNA samples were analysed, libraries were prepared using the TruSeq kit (Illumina), and sequenced on the NextSeq 550 platform as per manufacturer’s protocols.

### RNA-Seq analysis

#### Mapping reads and quantification

Coding regions of DNA and non-coding RNA from the mouse genome build 38 (mm10), downloaded from the Ensembl database, was used to create the transcriptome index. `kallisto` (Bray et al., 2016) was then used to pseudo-align RNA-seq reads for each library to these regions and quantify transcript abundance.

#### Differential expression analysis

Differential expression analysis between time points and expression normalization was performed using `DESeq2` (Love et al., 2014). Specifically, the `kallisto` abundance estimates were used as inputs to DESeq2, using a design matrix of timepoints. The `estimateSizeFactors` function was used to compute sizefactors for normalization, and the lfcShrink function, with `type=‘ashr’` for log-fold-change shrinkage (Stephens, 2016), was used to contrast timepoints. To select differentially expressed genes we used an FDR threshold of 0.001 and a log_2_-fold-change threshold of 1.5. We also use `DEseq2` to select a set of “house-keeping” or stably-expressed genes, by requiring these genes to display absolute log_2_ fold-changes < 0.1 for every successive day comparison, and have a mean (across days) normalized expression > 30 (this procedure selects 190 such stably-expressed genes).

#### Clustering

To cluster gene expression trajectories we take a Gaussian-mixture model (GMM) approach applied to *z*-transformed log_2_-values of normalized gene expression, for differentially expressed genes across the timepoints: day2-to-day3, day3-to-day4, day4-to-day5. In detail, we use the GMM implementation available in the Python `scikit-learn` library (Pedregosa et al., 2011), fitting GMMs on the trajectories for 20 random initializations. The number of clusters that best describe the data was determined using the minimum value of the average Bayesian Information Criterion (BIC) across random seeds, and the cluster allocation of each gene was done using consensusclustering, using the `ClusterEnsembles Python` library.

#### Gene ontology enrichment analysis

We applied a GO enrichment analysis on the three sets of genes corresponding to the broad patterns discovered using clustering, using the `Metascape` online tool (Zhou et al., 2019), https://metascape.org.

### ATAC-Seq Assays

On days 0–5 of optic vesicle differentiation, cells from 48 organoids were pooled and sorted using FACS as described above for the RNA-seq. One replicate was generated for each timepoint. ATAC-seq sample preparation was performed as described in Buenrostro et al. 2015. Briefly, 50,000 cells were lysed, and the transposition reaction carried out using the Tn5 transposase enzyme. PCR amplification of samples was conducted using a universal forward primer and a unique barcoded reverse primer for each time point. Samples were then purified and size selected. Bioanalysis showed that the libraries contained the expected fragment size distribution (Buenrostro et al., 2013). ATAC-seq samples were paired-end sequenced on the Illumina Hi-seq platform, with day 4 samples run separately from the other time points.

### ATAC-Seq analysis

#### Mapping reads, quantification and peak-calling

We used the `FastQC` tool (Andrews, 2010) and `cutadapt` (Martin, 2011) to perform quality control on and trim adapter sequences from the raw sequencing reads for all samples. Then we used bowtie2 (Langmead and Salzberg, 2012) to align trimmed reads to the mm10 genome. Duplicate reads, unmapped reads, and reads mapping to the mitochondrial genome were removed using `samtools` (Danecek et al., 2021). Filtered reads were shifted +4bp for the positive strand and −5bp for the negative strand to show the centre of the transposition site. bedtools (Quinlan and Hall, 2010) was used to generate the final bam and bed files used for downstream analysis. To enrich for fragments indicative of binding of TFs, we filter out reads corresponding to fragment sizes > 100bp (i.e. retain only sub-nucleosomal reads for downstream analysis). To identify genomic regions enriched for accessible chromatin we computationally call peaks using the `callpeak` function of `macs2` (Zhang et al., 2008), https://github.com/taoliu/MACS, with parameters `-f BAMPE -g mm -keep-dup all -B -SPMR -nomodel`, on paired-end bam files of each sample (day) separately. Using the narrowPeak `macs2` peak coordinates, we create a consensus set of peaks with which to study accessibility dynamics across days, using the `bedtools merge` command. From this consensus set of peaks we remove peaks overlapping blacklisted regions of the mm10 genome, as listed on http://mitra.stanford.edu/kundaje/akundaje/release/blacklists/. We use conditional quantile normalization (Hansen et al., 2012) to normalize signal in consensus ATAC-seq peaks while correcting for the effects of peak width and GC content. Due to the separate sequencing runs, day 4 samples showed considerable batch effects and were removed from subsequent analysis. Bedgraph files from macs2 were converted to BigWig files for visualisation in the UCSC genome browser. Finally we used `ChIPseeker` (Yu et al., 2015) to annotate peaks by their genomic context, collapsing peak annotations into four categories: promoter, intergenic, intronic and exonic.

#### Peaks by TAD regions

To assign peaks to TAD regions, we downloaded the coordinates for TAD regions from the work of Bonev et al. 2017 from the publicly-available https://github.com/aertslab/mucistarget/tree/master/data/tads. We note that there are often gaps between genomically adjacent TAD regions, and in these cases we take these inter-TAD regions to also define relevant regions to restrict the search-space of putative peak-gene links (interestingly, for the TAD data we use, the *Rax* and *Six6* genes are contained within such inter-TAD regions). EF-up, EF-down TADs are defined as those TADs containing the genes determined to be differentially up and down regulated across the day3-day5 transition. House-keeping TADs are defined as TAD regions containing genes that are stably expressed across the full timecourse.

#### GREAT analysis

To perform a first putative linking of dynamic peaks to genes we employed the online `GREAT` tool (McLean et al., 2010), http://great.stanford.edu to computationally generate region-gene associations. We use the coordinates of dynamic peak regions (regions of increasing and decreasing accessibility across day3-day5 transition) as `test regions`, and the whole genome as the `background regions` (default setting of GREAT).

#### Motif analysis of ATAC-seq dynamics

We obtain the DNA sequence of each consensus peak using `bedtools getfasta` and the mm10 genome. Following this we use `fimo` with parameters `-text -max-strand -thresh 1e-4` (i.e. a *p*-value threshold of 1e^−4^), we scan each peak region for occurrences of all motifs in the JASPAR2020 database (Mus Musculus TF motifs), using the presence/absence of motifs for further downstream analysis. We also restrict our attention to motifs of TFs that are *expressed* in the OV system, which we define to be that the mean (across RNA-seq replicates) `DEseq2`-normalized counts of a TF’s expression is > 50, at any point in the time course. To compute enrichment of motifs for our defined signal peak sets (peaks with increasing or decreasing accessibility signal above a log_2_ (f.c) of 1.5), we employ the Hypergeometric test (Fisher Test) using the set of peaks in TADs of house-keeping genes as a background.

Finally, to quantify the importance of each motif in a multivariate modelling approach, we fit a *L*2-regularised logistic regression model to predict the binary behaviour of each peak (increasing or decreasing accessibility signal) using the presence of motifs corresponding to EF-up, EF-down genes, day2-day3 differentially expressed genes as well as Sox2 and Otx2, as input features. Where there are multiple motifs for a TF, we collapse the input feature into a single feature in which a TF-motif is deemed to be present in a peak if that peak contains a significant occurrence of any of its possible motifs. We train models on two sets of peaks: dynamic peaks (those with log_2_(fc) > 1) within TADs containing EF-up genes and genome-wide dynamic peaks. We first use 5-fold cross-validation to determine an appropriate regularization parameter for each model, and then train models using these optimal parameters on the full sets of peaks. In this second step, we fit 10 models with different random initial seeds, and take the mean for each coefficient across each run. To perform this modelling we have used the `LogisticRegression` functions implemented in the `sklearn Python` library (Pedregosa et al., 2011).

#### Footprinting analysis

To quantify differences in putative TF binding between timepoints we used the TOBIAS framework (Bentsen et al., 2020). We first we used the bam files (with sub-nucleosomal reads) from each time point (days 0, 1, 2, 3 & 5) as input to `TOBIAS-ATACorrect` to correct for Tn5-insertion bias, resulting in ‘corrected’ bigwig files of ATAC-seq signal. Together with the coordinates for the consensus peak set, the latter were used as input to `TOBIAS-ScoreBigwig` to compute base-pair footprint scores for these genomic regions, again in bigwig format. Finally, these footprint bigwig files were was as input to `TOBIAS-BINDetect` (in time-series mode, using the `-time-series flag`) to identify motif occurrences in the consensus peak set and compute associated footprint scores for each day, for all mouse motifs in the JASPAR2020 database (Fornes et al., 2020). This final step also quantified differences in putative binding of TFs across the full consensus peak set between each pair of successive days (used to produce the results of Fig.4A). To generate the data for aggregate footprints (used for Fig.4B and Fig.S4.B), we used `TOBIAS-PlotAggregate` to average corrected ATAC-seq signal in ±100bp windows around occurrences of a given motif.

To quantify changes in aggregate footprints for a given TF-motif, we first compute two metrics, *flanking accessibility* and *footprint depth* (closely following the approach of Baek et al.2017), on the aggregate signal at each time point. For each motif we take the ±12bp window around the motif center as the ‘central region’ (putative region of binding), and firstly compute flanking accessibility as the mean aggregate signal in the 60bp either side of the central region. Footprint depth is then computed as the mean aggregate signal within the 24bp central region minus the flanking accessibility. Finally, we compute the difference between time points of each of these metrics to obtain the change in flanking accessibility and of footprint depth.

### CRISPR-Cas9 mediated disruption of candidate CREs

Mutations in the candidate regulatory elements within the Rax and Six6 TADs were introduced using CRISPR-Cas9. Guide RNAs, targeting the sites of TF binding motifs within the identified peaks, were individually cloned into the pSpCas9(BB)-2A-GFP (PX458, Addgene) vector. Rx-GFP cells were transfected with the resulting plasmid using Lipofectamine 3000 and cells selected based on GFP expression from the plasmid 48 hours later. Cells were plated at low density, individual clones were expanded and resulting cell lines were assessed for deletions over the CREs in question using Sanger sequencing. One cell line with a deletion encompassing motifs of interest along with one cell line without any deletions were maintained and used for optic vesicle organoid culture for each of the two selected putative CREs. At day 5, 24 organoids were pooled, and RNA isolated as described for the RNA-seq assay. The NanoString nCounter analysis system (Geiss et al.,2008) was used to quantify RNA expression for a panel of 180 genes whose expression levels change through the timecourse of differentiation.

#### Plotting and Visualization

Plots and visualizations were produced using the `matplotlib Python` (Hunter, 2007) library and `Inkscape` (Inkscape Project, 2020).

## Acknowledgements

We are grateful to Elisabeth Freyer and the IGC FACS facility, to the staff of the IGC Imaging facility for imaging support, to the IGC technical services team, to the genetics core at ECRF for sequencing and Alison Munro within the HTPU at the IGC for running the NanoString. We also thank Wendy Bickmore for her feedback on the manuscript.

## Contribution

L.J.O. performed the majority of the experimental work. J.R. contributed to the CRISPR-Cas9 mediated introduction of deletions and growing up of cell lines. H.B. contributed to the the RNA-seq and ATAC-seq assays. F.K. was responsible for initially setting up the organoid model system in our lab. A.S.P. designed and performed the computational analysis, with contributions from L.J.O.. A.S.P. and L.J.O wrote the manuscript. D.R.F. conceived the project and designed the experiments with contributions from L.J.O, H.B. and A.S.P..

## Data availability

Raw sequencing data generated in this work will be made publicly-available on GEO upon journal publication. We have explored Sox2 and Pou5f1 ChIP-seq and ChIP-nexus datasets publicly-available from GEO (accession codes GSE174774 and GSM4072776, GSM4072777). Analysis scripts and codes, as well as processed data are available at https://github.com/andrewpapa/optic_vesicle_eye_field_analysis.

