## Supplementary material for "Characterization of an Eye Field-like State during Optic Vesicle Organoid Development"

#### mESC Culture

Rx-GFP K/I EB5 mESCs (Wataya et al., 2008) obtained from the Riken BRC Cell Bank (cell number AES0145) were maintained under feeder-free conditions. Specifically, cells were cultured on 0.1% gelatin (Sigma, G2500) coated tissue culture flasks (Corning) in GMEM media (Gibco, 21710025) supplemented with 1X non-essential amino acids (Sigma, M7145), 1 mM Sodium pyruvate (Sigma, S8636), 10 $\mu$M 2-Mercaptoethanol (Gibco, 21985-023), 10% knock out serum replacement (KOSR) (Gibco, 10828-028), 1% Foetal Calf Serum (FCS) (Sigma, 12103C) and 500 U/ml leukaemia inhibitory factor (LIF) (produced in house). Care was taken to avoid over-confluency or undesirable differentiation of cells.

#### Optic Vesicle Organoid Culture

For the culture of optic vesicle organoids using a modified SFEBq (serum-free floating culture of embryoid body-like aggregates with quick reaggregation) technique, mouse ES cells were dissociated to single cells in TryplE and reaggregated in a differentiation media at a concentration of 4500 cells per 100 $\mu$l per well of a Nunclon Sphera 96-well U-bottomed low cell adhesion plate (Thermo Fisher Scientific, 174929). Two differentiation medias were used for this work due to the differentiation becoming unstable after the initial experiments had been performed. The organoids used for RNA and ATAC-seq were grown in CDM media and the organoids with mutations introduced in potential CREs were grown in KSR media. KSR differentiation media consisted of GMEM supplemented with 0.1 mM non-essential amino acids, 1 mM Sodium pyruvate, 10 $\mu$M 2-Mercaptoethanol and 1.5% KOSR. CDM differentiation media was made up of Iscove's Modified Dulbecco's Medium (IMDM) GlutaMAX$^{TM}$ Supplement (Thermo, 31980022) and Ham's F12 Nutrient Mix, GlutaMAX$^{TM}$ Supplement (Thermo, 31765027) mixed in a one-to-one ratio and supplemented with 1X Chemically Defined Lipid Concentrate (Thermo, 11905031), 5 mg/ml Bovine Serum Albumin (BSA) (Sigma, A3156-5G), 15 mg/ml bovine Apo-transferrin (Sigma, T1428) and 450 $\mu$M 1-Thioglycerol (Sigma, M6145). The day on which cells are aggregated and differentiation started was defined as day 0. For day 0 samples, cells were seeded in the low cell adhesion plates in stem cell maintenance media and collected after 24 hours (Fig.S1A). These cells were grown up to day 8 to ensure there was no differentiation or GFP expression (Fig.S1B).

[1]MRC Human Genetics Unit, Institute of Genetics and Cancer, University of Edinburgh, Edinburgh, UK
[2]School of Informatics, University of Edinburgh, Edinburgh, UK

On day 1, growth factor reduced Matrigel basement membrane matrix (Corning, 354230) was added to a final concentration of 2% (v/v). Cells were then incubated at 37°C with 5% CO2 and the aggregated cells differentiated to form optic vesicle like structures expressing Rx-GFP by day 5.

#### RNA Sequencing

**RNA Extraction, Quantification and Sequencing.** 24 organoids were pooled, washed with PBS 3 times or until all Matrigel and media was removed, and then dissociated to single cells in TryplE. Once dissociated, cells were resuspended in FACS buffer (PBS supplemented with 5% (vol/vol) FCS). Cells were sorted on the BD FACS Aria cell sorter, with cells gated manually into GFP positive and negative cell populations and collected in media. Cells were pelleted at 1200 RPM for 4 minutes at 4°C the supernatant was removed without disturbing the cell pellet and then cells were resuspended in 150 $\mu$l trizol. The Zymo Direct-zol RNA MicroPrep kit was used to prepare RNA samples according to the manufacturer's instructions including the optional DNaseI treatment. Following RNA extraction, samples were sent to the Wellcome Trust Clinical Research Facility at the Western General Hospital for quality and integrity analysis on the Agilent 2100 Bioanalyser using the RNA 6000 Nano chip. Samples were required to have an RNA Integrity Number of greater than 8. Concentration was quantified using the Qubit RNA broad range assay kit according to instructions. Illumina mRNA-seq libraries were prepared from 200 ng of total RNA using the TruSeq library prep kit. Libraries were pooled and sent for 75bp paired-end sequencing on the NextSeq 550 platform to generate around 40M reads per sample.

#### ATAC-seq

**Cell Lysis and Transposition Reaction.** Cells from 48 organoids were pooled and prepared for FACS as described above, with the cells counted and sorted into GFP expressing and non-expressing populations where possible. ATAC-seq sample preparation was performed as described in Buenrostro et al. 2015, with minor modifications. Cells were pelleted at 1200 RPM for 4 minutes at 4°C and then resuspended in ice cold PBS at a concentration of 100,000 cells per ml. 500 $\mu$l of cell suspension was pelleted, resuspended in 100 $\mu$l and centrifuged again before the supernatant was removed. Cells were then resuspended in 100 $\mu$l of cold ATAC lysis buffer (10 mM TrisCl, pH 7.4, 10 mM NaCl, 3 mM MgCl2, 0.1% IGEPAL CA-630) and kept on ice for 15 minutes, with occasional gentle pipetting. Samples were pelleted at 1200 RPM for 5 minutes at 4°C, supernatant discarded and resuspended in transposition mix consisting of 50 $\mu$l ChIPmentation buffer (10 mM Tris pH8, 5 mM MgCl2, 10% Dimethylformamide) and 2.5 $\mu$l Tn5 transposase enzyme (Illumina, 15027866) per sample. The transposition reaction was carried out at 37°C for 30 minutes. Samples were then purified using the Qiagen MinElute Reaction Cleanup Kit as per manufacturer's instructions, eluting in 10 $\mu$l elution buffer.

**PCR Amplification.** PCR amplification was conducted in a 32 $\mu$l reaction volume for each sample, made up of 10 $\mu$l transposed DNA, 4.7 $\mu$l of 10 $\mu$M custom barcoded nextera primer mix consisting of universal forward primer and one of the barcoded reverse primers, 1.3 $\mu$l 50X SYBR Green (Invitrogen, S7563) and 16 $\mu$l NEB-Next 2x PCR master mix (NEB, M0541S). The barcoded primers are detailed in Table S2. Samples were cycled as below:

1. 72°C for 5 min.
2. 98°C for 30 sec.
3. 98°C for 10 sec.
4. 63°C for 30 sec.
5. 72°C for 1 min.
6. Repeat steps 3-5 for 12 cycles.

Following PCR amplification, 30 $\mu$l of PCR reaction was made up to 50 $\mu$l with nuclease free $H_2O$.

**Size Selection.** To select against DNA over 1 Kb and remove primers, samples were purified using AMPure XP (Beckman Coulter, A63880) bead size selection. SPRI beads were brought to room temperature and vortexed and 50 $\mu$l added to each sample at a 1X ratio. Samples were incubated at room temperature for 5 minutes to allow fragment binding. Reactions were then placed in a magnetic stand and allowed to separate for 5 minutes, the supernatant discarded, and the beads washed twice with 200 $\mu$l 80% ethanol for 30 seconds. The samples were left to air dry for 10 minutes. 20 $\mu$l 1X TE buffer was added for 30 seconds to elute DNA from the beads. Samples were returned to the magnetic stand to separate the beads from the samples.

**Quantification and Sequencing.** Before sequencing, the quality and quantity of samples was checked. The total amount of DNA was measured using the Qubit dsDNS high-sensitivity assay as per manufacturer's instructions. For quality assessment ATAC-seq samples underwent high-sensitivity DNA bioanalysis at Edinburgh Genomics. Bioanalysis showed that the libraries contained the expected fragment size distribution, containing peaks corresponding to mononucleosomal, dinucleosomal and trinucleosomal fragments as described in the published protocol (Buenrostro et al., 2013). ATAC-seq samples were sent for 75bp paired-end sequencing on the Illumina Hi-seq platform at Edinburgh Genomics. Day 4 samples were sequenced on a separate run to the other time points resulting in the batch effects that led to this time point being excluded from much of the analysis.

### Genome Editing in mESCs

**CRISPR-Cas9 Mutant Cell Line Generation.** CRE null mESCs and their WT counterparts were generated, using CRISPR-Cas9, from the Rx-GFP cell line. CRISPR single guide RNAs were designed targeting the site of TF binding sequences for the chosen *Rax* and *Six6* peaks. Sequences of the guide RNAs used are detailed in Table S3.

The pSpCas9(BB)-2A-GFP (PX458) (Addgene, 48138) plasmid vector (Ran et al, 2013) was linearized by digestion with BbsI, gel purified and ligated with an annealed pair of guide oligos. The resulting plasmid DNA was transformed into chemically competent DH5$\alpha$ cells and purified from liquid culture using the QIAprep Spin Miniprep Kit as per manufacturer's instructions.

Rx-GFP mESCs were transfected with 2 $\mu$g of plasmid DNA diluted in Opti-MEM (ThermoFisher, 31985062) and 6 $\mu$l Lipofectamine 2000 transfection reagent (ThermoFisher, 11668019). Media was changed after 6 hours. After 48 hours transfected cells were sorted using FACS based on GFP expression. Cells were plated at a density of 1000 cells per 10 cm dish and grown for around 10 days or until colonies began to appear. Colonies were picked and plated in duplicate in a 96-well plate. One well was used to extract genomic DNA from the cells. The region targeted by the guide RNA was amplified and Sanger sequenced using primers as detailed in Table S4. Initially, cell lines were genotyped using the primers closest to the guide cut site.

Cell lines with deletions encompassing the targeted TFBSs and control clones, that had no detectable mutations introduced, were expanded from the duplicate 96-well plate. Once expanded, the appropriate distal primers were used to amplify a larger region around the guide cut site and Sanger sequenced, both to confirm the deletions are present in the expanded cell line and to check for any larger heterozygous deletions that may have been missed by using the initial primers.

**Nanostring nCounter RNA Quantification.** A custom NanoString CodeSet of 200 genes was designed to include genes that were differentially expressed for each sequential timepoint comparison, including genes that were up and down regulated and 20 housekeeping genes. 24 organoids were pooled, and RNA was extracted as described for the RNA-seq assay. The NanoString nCounter analysis was performed by the HTPU within the IGC. For the hybridisation reactions, 70 $\mu$l of Hybridisation Buffer was added to each vial of the Reporter CodeSet, 8 $\mu$l of this was added to each of the hybridisation tubes. 5 $\mu$l of 20 ng/$\mu$l RNA (100 ng total) was added to the appropriate hybridisation tube, followed by 2 $\mu$l of the Capture Probeset. Tubes were incubated at 65°C for 18 hours. Following hybridisation, the samples were processed using the nCounter Prep station within 24 hours of hybridisation. The hybridised RNA samples and all components of the nCounter masterkit were loaded in the prep station and processed using the high sensitivity protocol. The prep station was used to purify the hybridised samples by removing excess probes, then binding, immobilising and aligning them in a sample cartridge for analysis. At this point, each colour-coded barcode is attached to a single target-specific probe corresponding to an analyte of interest. The cartridges were sealed and read in the digital analyser using the max setting to count 555 FOV (Field of View). The Reporter code counts for each sample, as produced by the Digital Analyser, were QCed and normalised using a combination of positive control targets and CodeSet content normalisation, which uses housekeeping genes, to apply a sample specific correction factor to all target probes within that sample lane.

| Organoid development day | Total Live Cells | Total GFP+ cells | % GFP+ |
|---|---|---|---|
| day5 (n=35) | 12,731 | 9,404 | 74 |
| day6 (n=39) | 19,313 | 13,566 | 72 |
| day7 (n=38) | 19,013 | 12,002 | 65 |
| day8 (n=15) | 17,811 | 9,752 | 54 |

**Table S1. Counts of live cells from single dissociated organoids and associated percentages of GFP positive cells.**

| | |
|---|---|
| Ad1_noIndex | AATGATACGGCGACCACCGAGATCTACACTCGTCGGCAGCGTCAGATGTG |
| Ad2.1_TAAGGCGA | CAAGCAGAAGACGGCATACGAGATTCGCCTTAGTCTCGTGGGCTCGGAGATGT |
| Ad2.2_CGTACTAG | CAAGCAGAAGACGGCATACGAGATCTAGTACGGTCTCGTGGGCTCGGAGATGT |
| Ad2.3_AGGCAGAA | CAAGCAGAAGACGGCATACGAGATTTCTGCCTGTCTCGTGGGCTCGGAGATGT |
| Ad2.4_TCCTGAGC | CAAGCAGAAGACGGCATACGAGATGCTCAGGAGTCTCGTGGGCTCGGAGATGT |
| Ad2.5_GGACTCCT | CAAGCAGAAGACGGCATACGAGATAGGAGTCCGTCTCGTGGGCTCG |
| Ad2.6_TAGGCATG | CAAGCAGAAGACGGCATACGAGATCATGCCTAGTCTCGTGGGCTCGGAGATGT |
| Ad2.7_CTCTCTAC | CAAGCAGAAGACGGCATACGAGATGTAGAGAGGTCTCGTGGGCTCGGAGATGT |
| Ad2.8_CAGAGAGG | CAAGCAGAAGACGGCATACGAGATCCTCTCTGGTCTCGTGGGCTCGGAGATGT |

**Table S2. Nextera PCR primers used for ATAC-seq sample preparation.** Ad1 was the forward primer common to all samples, whereas barcoded primers 2.1-2.8 were unique to each sample.

| Guide Name | Guide RNA Sequence |
|---|---|
| Rax Forward | CACCG**TGTAGATTAGCTCCTAACAA** |
| Rax Reverse | AAAC**TTGTTAGGAGCTAATCTACA**C |
| Six6 Forward | CACCG**ATAATCTCTTTAATTGGTGT** |
| Six6 Reverse | AAAC**ACACCAATTAAAGAGATTAT**C |

**Table S3. Guide RNA sequences for disruption of *Rax* and *Six6* candidate *cis*-regulatory elements.**

| Primer Name | Primer Sequence |
|---|---|
| Rax TFBindingSite Fwd | TGGAGCCTGCCTTTGTGTAG |
| Rax TFBindingSite Rev | CAGGTTGGAGCTGGGAAGAG |
| Distal Rax Fwd | TCCCAGCTGGCTAGGTAGAG |
| Distal Rax Rev | GAAGCAGTGCATGCTGGATA |
| Six6 TFBindingSite Fwd | CACAGTGCCAACATGCAAGT |
| Six6 TFBindingSite Rev | AGCAGGCTTTCCAAAGAGGT |
| Distal Six6 Fwd | ACAGGAGGGAGACTATGATTGG |
| Distal Six6 Rev | AGTCGCATAAGACACCGTGG |

**Table S4. Sequencing primers used for amplification of putative *Rax* and *Six6 cis*-regulatory elements and Sangar sequencing for confirmation of mutations**

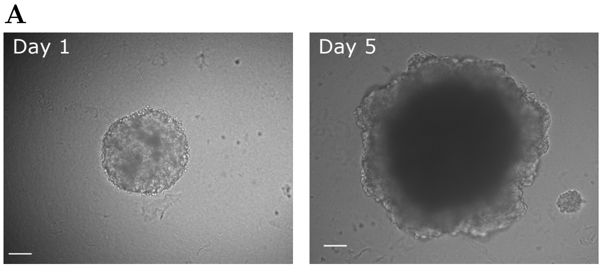


**Fig. S1. Organoid culture controls for OV timecourse.** Representative images of organoids cultured in maintenance media showing no GFP expression. Scale bars: 100 $\mu$m.

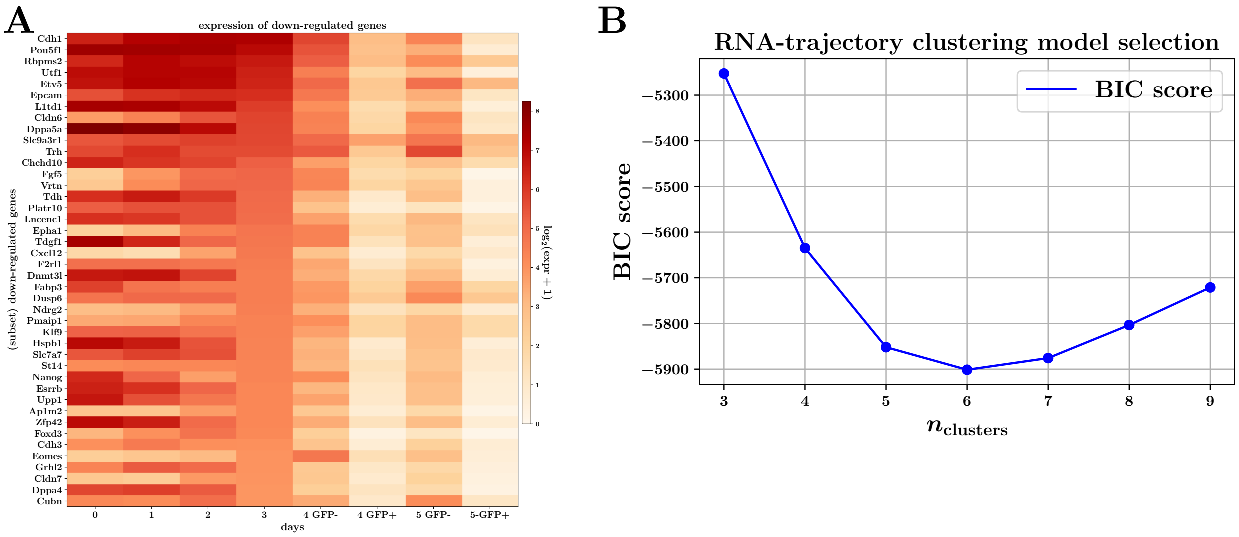


**Fig. S2. Down-regulated genes indicate pluripotency and other gene-expression programs are suppressed during eye field specification. A.** Expression heatmap displaying expression changes across the OV timecourse, for a selection of genes down-regulated across day3-day4/5 transition. **B.** BIC model selection plot for $n_{\text{clusters}}$ parameter in GMM-clustering.

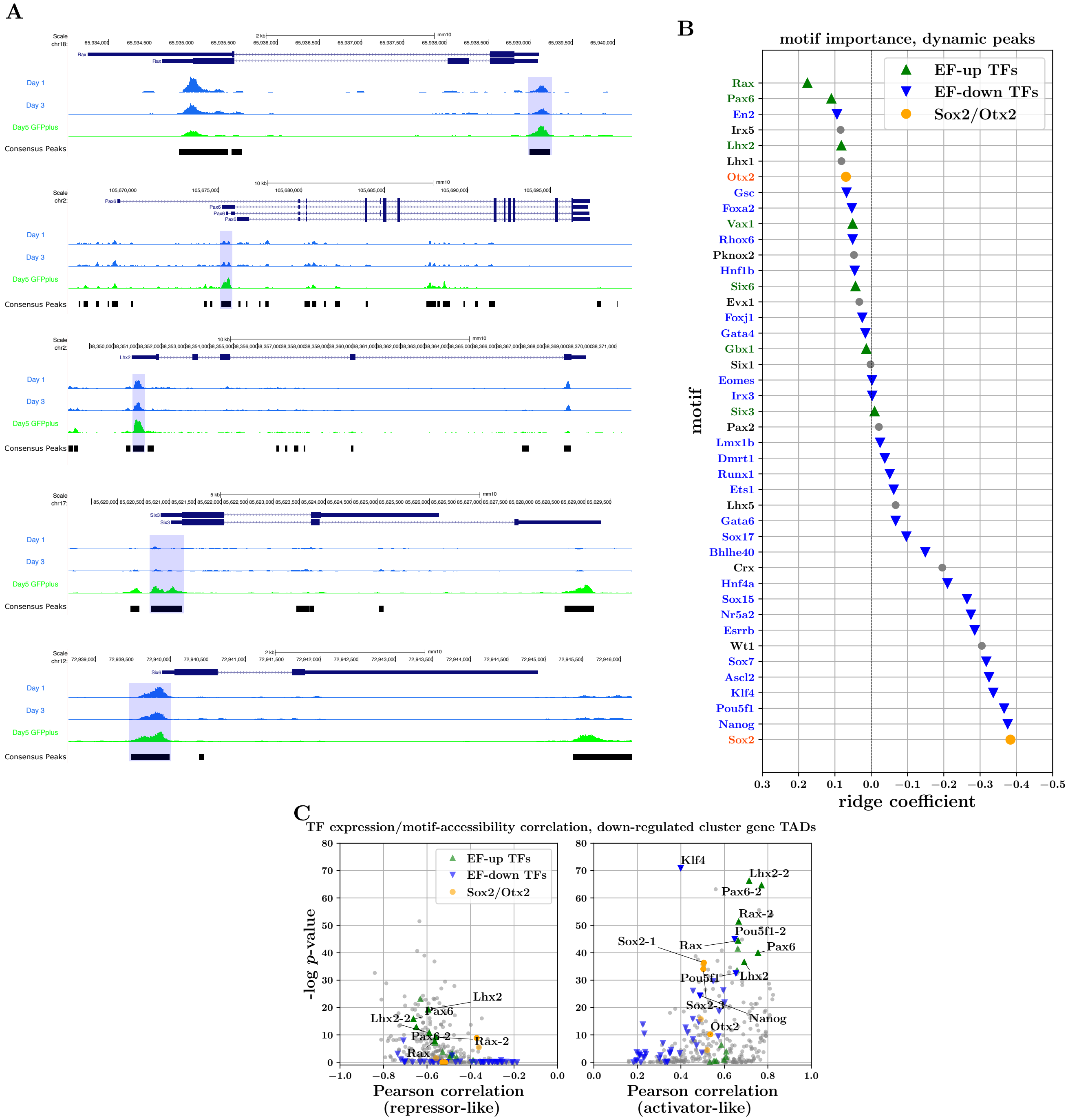


**Fig. S3.** **Promoters of canonical eye field TFs are accessible from early in OV timecourse. Motif importance for genome-wide dynamic peaks mirrors importance in EF TADs.** **A.** Bigwig tracks displaying changes in accessibility for regions around canonical EFTFs across the OV development timecourse. Promoters regions of these TFs have been highlighted in blue. **B.** Coefficients of logistic-regression model trained to predict opening-vs-closing behaviour of genome-wide dynamic ATAC-seq peaks, using presence of TF-motifs as input covariates. Magnitude of coefficients is indicative of importance of respective TF-motifs for these predictions. **C.** TF-expression/TF-motif-accessibility correlations for EF-down TAD peaks. RHS and LHS plots illustrate the median across positive and negative correlations respectively, indicative of activator-like and repressor-like behaviour of the respective TFs.

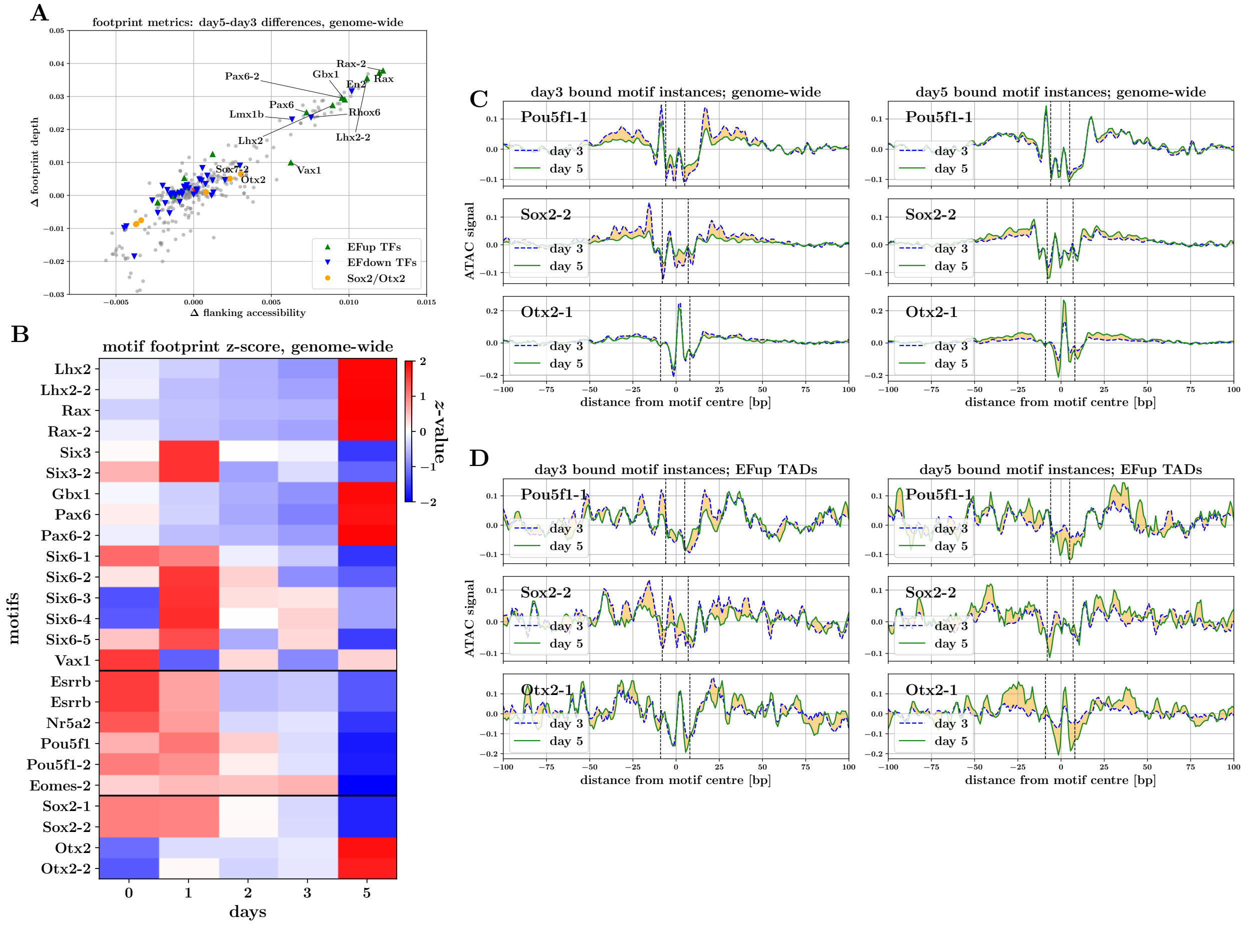


**Fig. S4. Footprinting patterns genome-wide. A.** Scatterplot of day3-day5 differences in TF-motif footprint depth versus flanking accessibility, for peaks genome-wide. **B.** Heatmap of TF-motif footprint scores in peaks genome-wide, $z$-transformed across time-course (median across peaks). **C. & D.** Aggregate footprint signals for Pou5f1, Sox2 and Otx2 binding motifs. Plots illustrate aggregate signal on day 3 and day 5 for motif-instances predicted to be bound on day 3 (left) and day 5 (right). **C.** Aggregate footprint signals for genome-wide motif instances. **D.** Aggregate footprint signals for EF-up TAD motif instances.

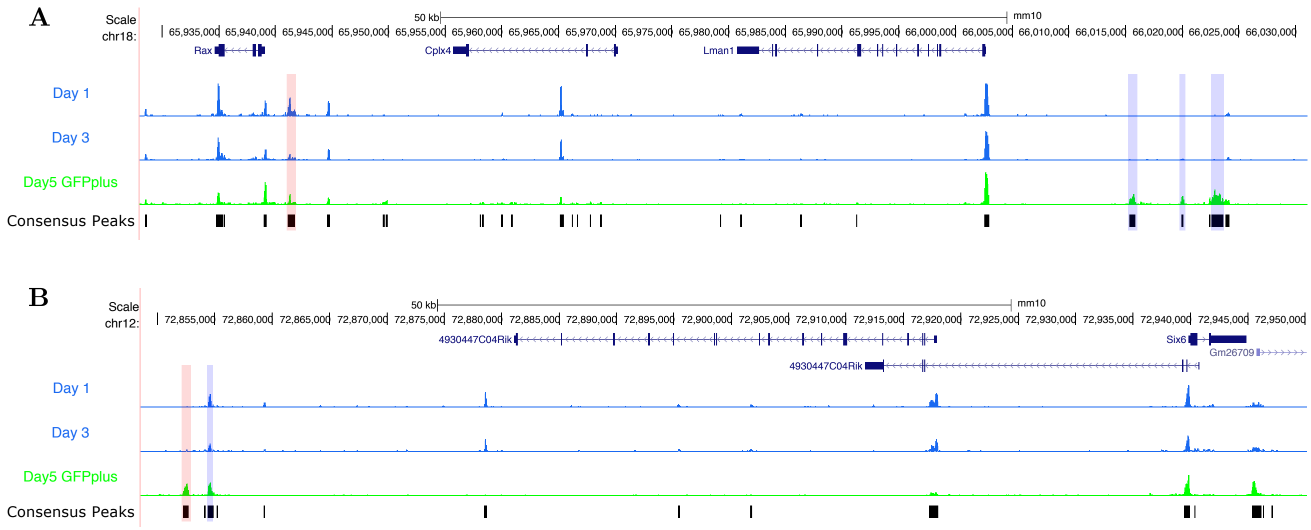


**Fig. S5. Accessibility changes around candidate *Rax* and *Six6* enhancer elements. A. & B.** Changes in ATAC-seq signal around the *Rax* and *Six6* loci, with consensus peaks illustrated by black bars. Peaks showing interesting footprint score changes mentioned in text are highlighted in blue, and the peaks chosen for perturbation in red.

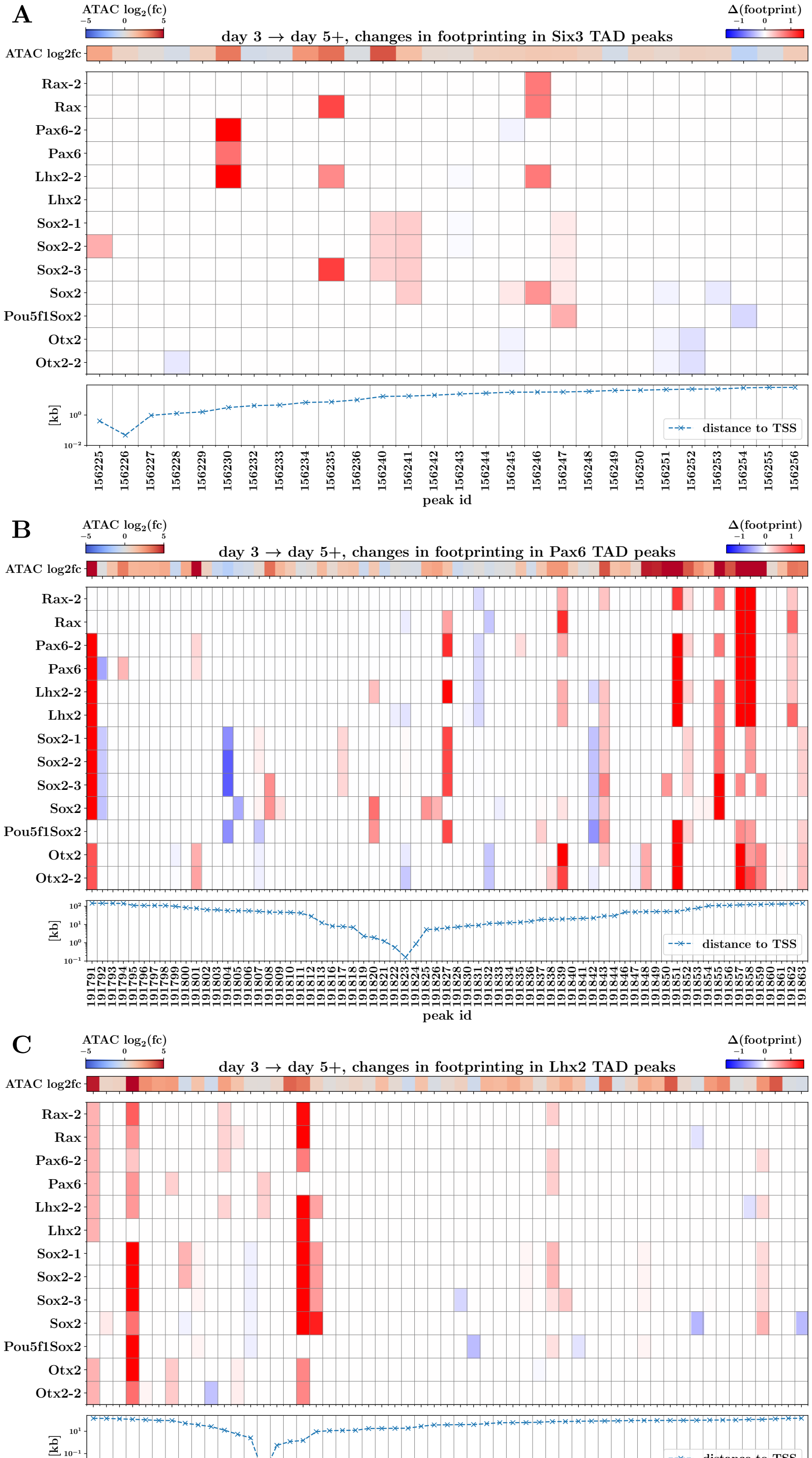


**Fig. S6. Changes in motif footprint scores of key TFs identify candidate *Pax6*, *Lhx2* and *Six3 cis*-regulatory elements. A.** Heatmap of day3-to-day5 changes in footprint scores for motifs of key TFs at peaks contained in *Six3* TAD region. Also illustrated are associated changes in normalized ATAC-seq signal for each peak (upper panel), and distance of each peak from the *Six3* TSS (lower panel). **B.** & **C.**: same as **A.** but displaying changes for peaks within *Pax6* and *Lhx2* TAD regions.

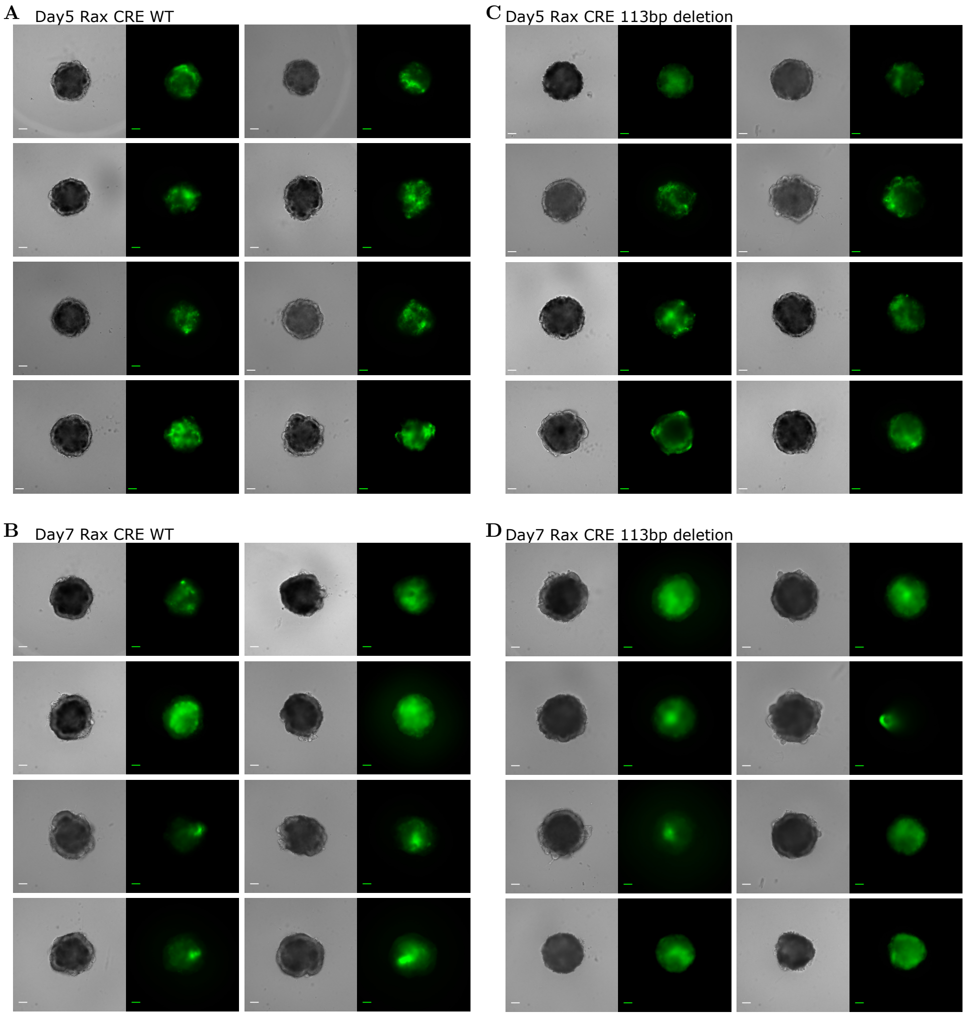


**Fig. S7.** **Organoids derived from candidate *Rax* enhancer CRISPR-disrupted cell lines. A. & B.** Organoids derived from wildtype *Rax* enhancer CRISPR cell-line, at days 5 and 7. **C. & D.** Organoids derived from mutant *Rax* enhancer CRISPR cell-line, at days 5 and 7. Scale bars: $100\ \mu$m.

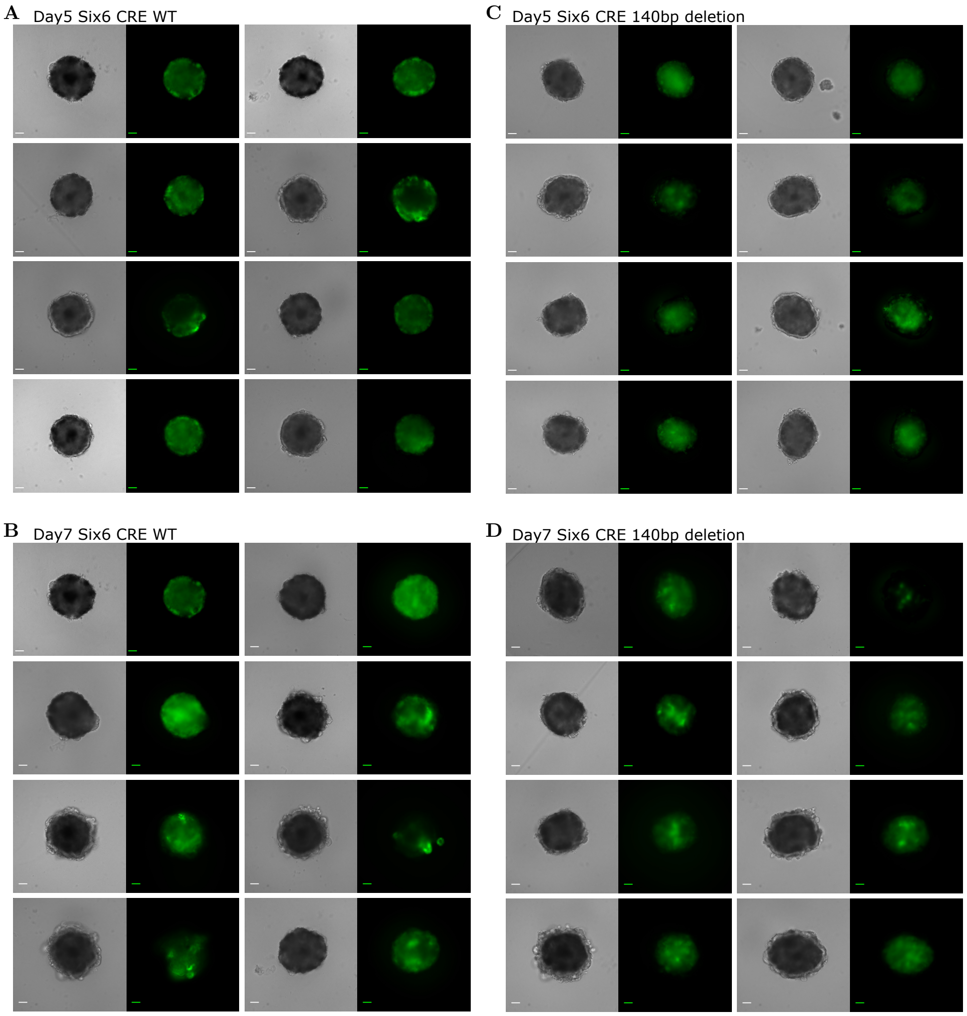


**Fig. S8. Organoids derived from candidate *Six6* enhancer CRISPR-disrupted cell lines. A. & B.** Organoids derived from wildtype *Six6* enhancer CRISPR cell-line, at days 5 and 7. **C. & D.** Organoids derived from mutant *Six6* enhancer CRISPR cell-line, at days 5 and 7. Scale bars: $100\ \mu$m.

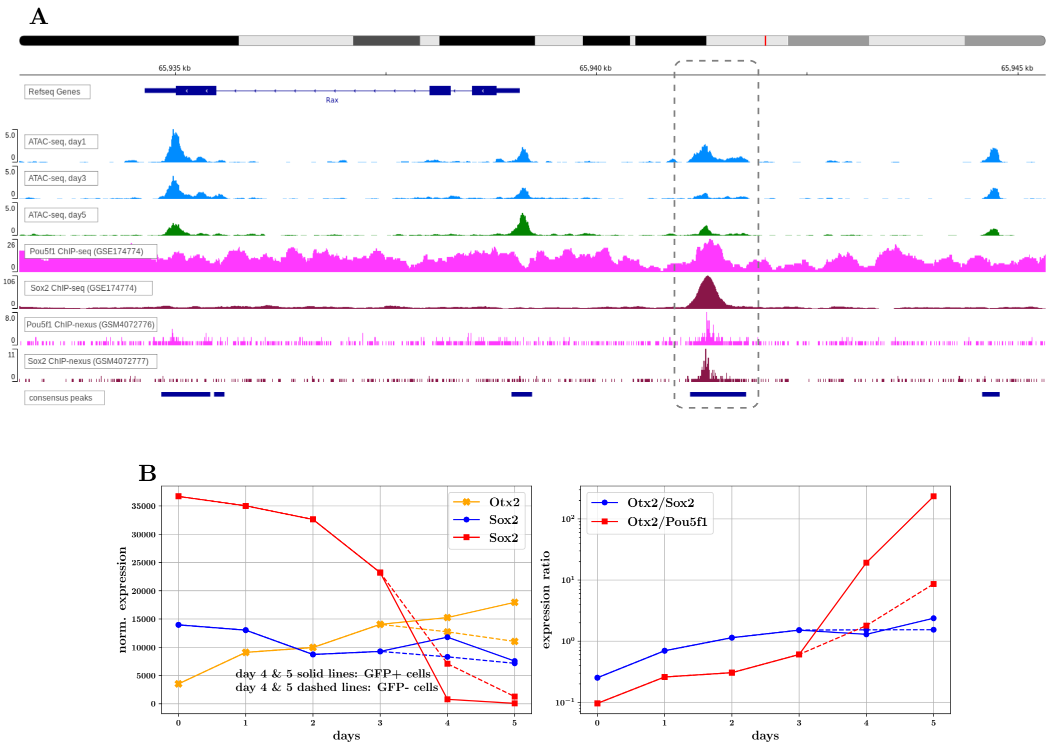


**Fig. S9.** **ATAC-seq, ChIP-seq and RNA-seq data lead to hypothesis of Rax element activation. A.** Bigwig tracks for day 1, 3 & 5 ATAC-seq signal (our organoid data) and E14 mESC Pou5f1 and Sox2 ChIP-seq and ChIP-nexus data around *Rax* locus. Candidate regulatory element highlighted in dashed box overlaps with putative *Rax* enhancer element identified by Danno et al. 2008 and perturbed in the OV-organoid system using CRISPR. **B.** Expression (left) and expression-ratios (right) for key TFs hypothesized to play a role in the activation of the proximal Rax element.
